# A Th17 cell-intrinsic glutathione/mitochondrial-IL-22 axis protects against intestinal inflammation

**DOI:** 10.1101/2023.07.06.547932

**Authors:** Lynn Bonetti, Veronika Horkova, Joseph Longworth, Luana Guerra, Henry Kurniawan, Davide G. Franchina, Leticia Soriano-Baguet, Melanie Grusdat, Sabine Spath, Eric Koncina, Anouk Ewen, Carole Binsfeld, Charlène Verschueren, Jean-Jacques Gérardy, Takumi Kobayashi, Catherine Dostert, Sophie Farinelle, Janika Härm, Ying Chen, Isaac S. Harris, Philipp A. Lang, Vasilis Vasiliou, Ari Waisman, Elisabeth Letellier, Burkhard Becher, Michel Mittelbronn, Dirk Brenner

## Abstract

Although the intestinal tract is a major site of reactive oxygen species (ROS) generation, the mechanisms by which antioxidant defense in gut T cells contribute to intestinal homeostasis are currently unknown. Here we show, using T cell-specific ablation of the catalytic subunit of glutamate cysteine ligase (*Gclc*), that the ensuing loss of glutathione (GSH) impairs the production of gut-protective IL-22 by Th17 cells within the lamina propria. Although *Gclc* ablation does not affect T cell cytokine secretion in the gut of mice at steady-state, infection with *C. rodentium* increases ROS, inhibits mitochondrial gene expression and mitochondrial function in *Gclc*-deficient Th17 cells. These mitochondrial deficits affect the PI3K/AKT/mTOR pathway, leading to reduced phosphorylation of the translation repressor 4E-BP1. As a consequence, the initiation of translation is restricted, resulting in decreased protein synthesis of IL-22. Loss of IL-22 results in poor bacterial clearance, enhanced intestinal damage, and high mortality. ROS-scavenging, reconstitution of IL-22 expression or IL-22 supplementation *in vivo* prevent the appearance of these pathologies. Our results demonstrate the existence of a previously unappreciated role for Th17 cell-intrinsic GSH coupling to promote mitochondrial function, IL-22 translation and signaling. These data reveal an axis that is essential for maintaining the integrity of the intestinal barrier and protecting it from damage caused by gastrointestinal infection.

**Executive summary:**

- GSH-regulated Th17 cell-derived IL-22, but not IL-17 is required to maintain intestinal barrier integrity and to revent lethality following *C. rodentium* infection.
- *GCLC* expression in IBD patients correlates positively with expression of genes related to gut integrity.
- *Gclc*-deficient Th17 cells accumulate mitochondrial ROS, which is linked to impaired mitochondrial function, ysregulated PI3K/AKT/mTOR signaling and impaired translation of IL-22.
- ROS-scavenging, IL-22 reconstitution or T cell-specific expression of IL-22 in *Gclc*-deficient T cells rescues utant mice from the lethal infection outcome *in vivo*.

## Introduction

T helper (Th) cells are essential for the protection of mucosal surfaces. In the gastrointestinal (GI) tract, Th cells and in particular Th17 cells, maintain gut homeostasis by inducing tolerance to the microbiome and by defending against intestinal pathogens (van Wijk and Cheroutre, 2010). Although Th17-derived cytokines such as IL-17 and IL-22 have been linked to inflammatory bowel disease (IBD), increasing evidence points to beneficial effects of these cytokines during intestinal inflammation (Eken et al., 2014, Kamanaka et al., 2011, Fujino et al., 2003, Brand et al., 2006, Monteleone et al., 2012). IL-22 and IL-17 induce antimicrobial peptide (AMP) production by intestinal epithelial cells (IECs) and protect the intestinal barrier against pathogens that directly efface the intestinal epithelium (Liang et al., 2006, Kim et al., 2012, Tsai et al., 2017, Keir et al., 2020, Lee et al., 2015). Accordingly, neutralization or genetic ablation of IL-22 results in severe disease and increased epithelial cell damage in mouse colitis models (Sugimoto et al., 2008, Zheng et al., 2008, Basu et al., 2012). In human IBD, damage to the epithelial layer and increased permeability have been proposed as primary defects (Shorter et al., 1972, Corridoni et al., 2014). Such intestinal damage is often causally linked to reactive oxygen species (ROS), in line with a striking reduction in intestinal antioxidant capacity and a rise in oxidative stress (Sido et al., 1998, Buffinton and Doe, 1995, Lih-Brody et al., 1996, Kruidenier et al., 2003, Aviello and Knaus, 2017). However, how antioxidant defense provided by gut T cells contributes to intestinal integrity is unclear.

The main antioxidant produced by activated T cells is glutathione (GSH) (Mak et al., 2017). The rate-limiting reaction of GSH synthesis is catalyzed by glutamate cysteine ligase (GCL), a complex of catalytic (GCLC) and modifier (GCLM) subunits (Meister, 1983, Chen et al., 2005). Here, we use a T cell-specific *Gclc*-deficient mouse model to show that a lack of *Gclc* impairs Th17 cell production of IL-22 in response to the intestinal pathogen *Citrobacter rodentium* (*C. rodentium*), leading to defective bacterial clearance, enhanced intestinal damage, and high mortality. Our results demonstrate that GSH regulates mitochondrial function and ROS in Th17 cells, which links the mitochondrial activity to the expression of IL-22 and is critical for combatting a GI infection.

## Results

### *Gclc* ablation in murine T cells results in high mortality during GI infection

*C. rodentium* causes colitis symptoms in mice and shares pathogenic mechanisms with human *E. coli* intestinal pathogens (Collins et al., 2014, Mullineaux-Sanders et al., 2019). We infected wild-type (WT) C57BL/6 mice with *C. rodentium* and isolated colonic lamina propria (LP) cells at day 7 post-infection (p.i.) in order to investigate antioxidant responses in these cells. Flow cytometric analysis revealed that CD4^+^ T cells of infected WT mice produced more thiols than those of uninfected WT mice (Figure 1A), indicating a greater antioxidant capacity. The heightened metabolic activity of activated T cells drives increased ROS production, which is in line with the expanded antioxidant capacity of T cells during *C. rodentium* infection (Gülow et al., 2005, Yi et al., 2006, Sena et al., 2013). We hypothesized that LP CD4^+^ T cells increase their production GSH, the main intracellular antioxidant and thiol, to neutralize the rising oxidative stress caused by infection. Indeed, intracellular GSH was increased in sorted LP CD4^+^ cells isolated from infected WT mice (Figure 1B). Cytosolic ROS levels were comparable in LP CD4^+^ cells isolated from uninfected and infected WT mice (Figure 1C), reflecting superior ROS scavenging by the elevated GSH in the infected animals.

**Figure 1:**
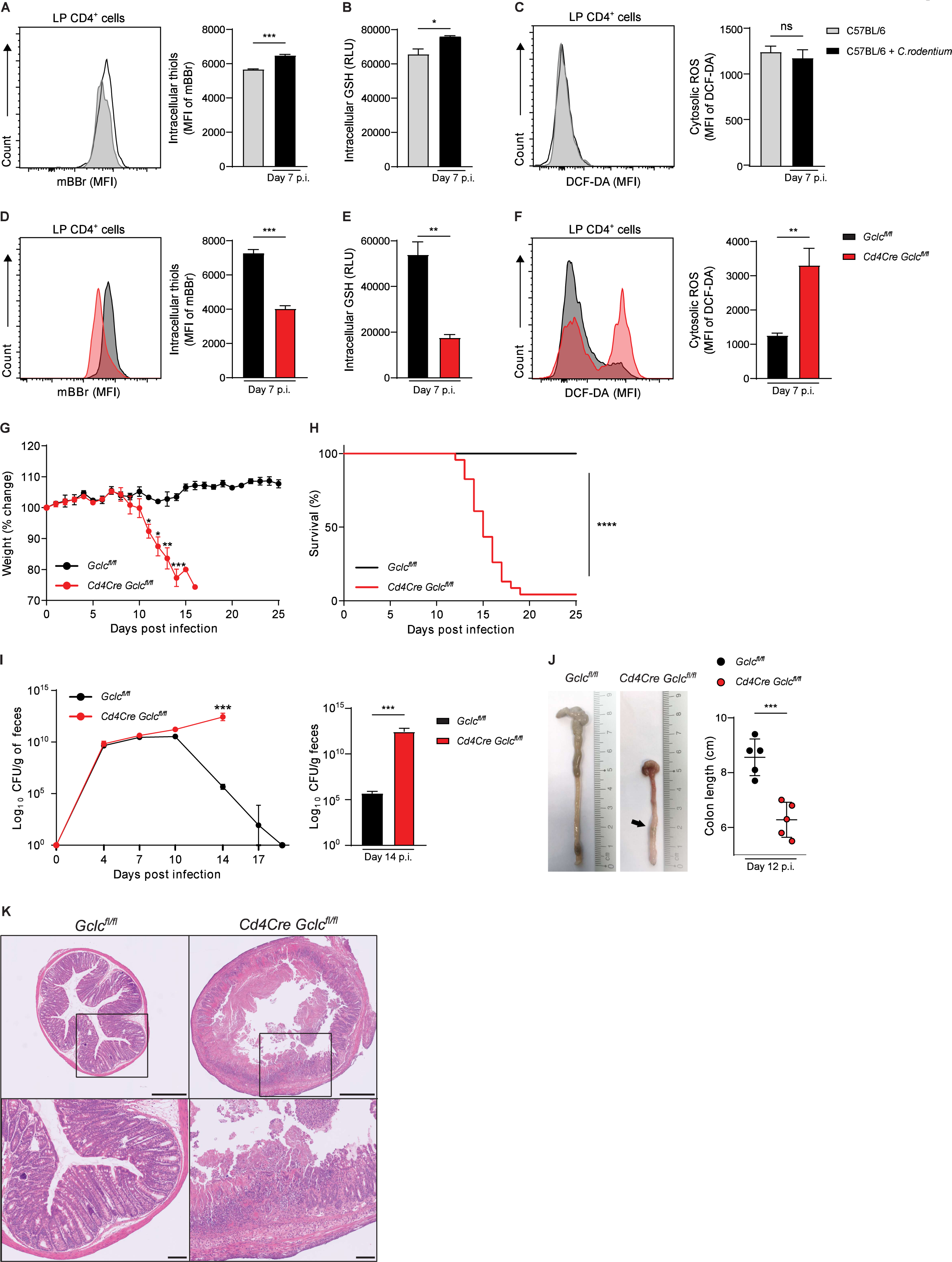
*Gclc* in LP T cells controls ROS and protects mice from lethal *C. rodentium* infection. (A) Left: Flow cytometric analysis (FCA) of intracellular thiol levels as detected by mBBr staining of CD4^+^ cells in colonic LP, which was isolated from uninfected and *C. rodentium*-infected *C57BL/6* mice at day 7 p.i. MFI, mean fluorescence intensity. Right: Quantification of left panel results. Data are mean±SEM (n=3-4). (B) Quantification of luminescence-based assay of intracellular GSH in FACS-sorted colonic LP CD4^+^ cells isolated as in (A). Data are mean±SEM of relative luminescence units (RLU) (n=3). (C) Left: FCA of intracellular ROS in CD4^+^ cells of the colonic LP, which was isolated as in (A) and subjected to DCF-DA staining. Right: Quantification of left panel results. Data are mean±SEM (n=3). (D) Left: FCA of intracellular thiol levels as detected by mBBr staining in CD4^+^ cells of the colonic LP, which was isolated from *C. rodentium*-infected *Gclc^fl/fl^* and *Cd4Cre Gclc^fl/fl^* mice at day 7 p.i. Right: Quantification of left panel results. Data are mean±SEM (n=3) and representative of 2 trials. (E) Quantification of luminescence-based assay of intracellular GSH in FACS-sorted CD4^+^ cells from colonic LP isolated as in (D). Data are mean±SEM of RLU (n=3). (F) Left: FCA of intracellular ROS in CD4^+^ cells of the colonic LP, which was isolated as in (D) and subjected to DCF-DA staining. Right: Quantification of left panel results. Data are mean±SEM (n=4). (G) Change in whole body weight of *Gclc^fl/fl^* and *Cd4Cre Gclc^fl/fl^* mice infected with *C. rodentium* on day 0 and assayed on the indicated days p.i. Data are mean±SEM (n=3-4) and representative of 5 trials. (H) Survival of *Gclc^fl/fl^* (n=14) and *Cd4Cre Gclc^fl/fl^* (n=23) mice infected with *C. rodentium* on day 0. Data are pooled from 5 independent trials. (I) Left: CFU of *C. rodentium* in feces of *Gclc^fl/fl^* and *Cd4Cre Gclc^fl/fl^* mice at the indicated days p.i. Data are mean±SEM (n=3-4) and representative of 5 trials. Right: Statistical analysis of left panel data at day 14 p.i. (J) Left: Macroscopic views and lengths of colons isolated from *C. rodentium*-infected *Gclc^fl/fl^* and *Cd4Cre Gclc^fl/fl^* mice at day 12 p.i. Right: Quantification of colon lengths from left panel results. Data are mean±SEM (n=5); 2 trials. (K) Top: H&E-stained sections of distal colon from *C. rodentium*-infected *Gclc^fl/fl^* and *Cd4Cre Gclc^fl/fl^* mice at day 12 p.i. Data are representative of 3 mice/genotype. Scale bars, 500µm. Bottom: Higher magnification views of the boxed areas in the top panels. Scale bars, 100µm.

These data prompted us to investigate whether GSH is necessary to buffer ROS in LP T cells of *C. rodentium*-infected mice. We took advantage of a mutant mouse strain harboring a T cell-specific deletion of *Gclc* (*Cd4Cre Gclc^fl/fl^* mice); T cells of these mutants cannot synthesize GSH (Mak et al., 2017). As expected, we observed significantly lower levels of intracellular thiols and GSH in LP CD4^+^ cells of infected *Cd4Cre Gclc^fl/fl^* mice compared to those of infected *Gclc^fl/fl^* littermate controls (Figure 1D, 1E). Consequently, cytosolic ROS levels were increased in LP CD4^+^ cells of infected mutants compared to controls (Figure 1F), confirming that GSH loss reduces ROS buffering capacity during infection. Interestingly, cytosolic ROS in T cells of uninfected *Cd4Cre Gclc^fl/fl^* and *Gclc^fl/fl^* mice were comparable (Figure S1A), consistent with our previous finding that higher ROS occur in *Gclc*-deficient splenic CD4^+^ T cells only after activation (Mak et al., 2017). Thus, GSH is not needed to control ROS in T cells at steady-state but becomes crucial upon infection.

To investigate physiological consequences of GSH loss in T cells, we monitored weights of littermate *Cd4Cre Gclc^fl/fl^* and *Gclc^fl/fl^* mice following *C. rodentium* infection. *Cd4Cre Gclc^fl/fl^* mice showed significant weight loss by day 11 p.i., whereas control animals were unaffected (Figure 1G). Over 95% of mutant mice succumbed to the infection by day 20 p.i., whereas no mortality was observed among controls (Figure 1H). No weight loss or spontaneous disease development was observed in uninfected mutant mice up to age 12 months (data not shown). While bacterial clearance, as determined by measuring *C. rodentium* load in feces, was achieved by day 20 p.i. in control mice, the mutants showed a steady increase in bacterial load until death (Figure 1I, left). At day 14 p.i., fecal *C. rodentium* shedding in mutant mice was increased by >10^5^-fold relative to controls (Figure 1I, right). Anatomically, infected *Cd4Cre Gclc^fl/fl^* mice displayed a marked shortening of colon length on day 12 p.i. (Figure 1J). Moreover, these colons exhibited ulcers and thickening that were not visible on colons from infected *Gclc^fl/fl^* mice (Figure 1J, S1B). Intestinal crypt loss, ulceration and necrosis of the intestinal epithelium were present in distal colons of infected *Cd4Cre Gclc^fl/fl^* mice but not in those of infected controls (Figure 1K). Of note, intestinal microbiota composition was comparable in uninfected *Cd4Cre Gclc^fl/fl^* mice and controls (Figure S1C). Thus, *Gclc* ablation in T cells renders mice unable to control *C. rodentium* infections, leading to intestinal damage and high mortality. These data suggest a protective role for T cell-derived GSH during GI infection.

### Pathogenic Th17 cells are the dominant Th subset responding to *C. rodentium*

*C. rodentium* infection elicits a robust T cell response in mice that ensures bacterial clearance and survival (Simmons et al., 2003, Bry and Brenner, 2004, Bry et al., 2006). Accordingly, we observed higher frequencies and total numbers of activated/memory CD4^+^ T cells (CD62L^-^ CD44^+^), and reduced frequencies of naïve CD4^+^ T cells (CD62L^+^CD44^-^), in colonic LP isolated from infected *Gclc^fl/fl^* mice at day 7 p.i. compared to uninfected controls (Figure S2A, S2B). To determine which T cell subsets contribute to this response, we analyzed cytokine production in colonic CD4^+^ T cells from infected and uninfected *Gclc^fl/fl^* mice by flow cytometry. We found comparable frequencies of Th22 cells (CD4^+^IL-17^-^IL-22^+^) and conventional Th17 cells producing IL-17 but not IL-22 (CD4^+^IL-17^+^IL-22^-^) (Figure 2A-C). However, the frequency of so-called “pathogenic” Th17 cells producing both IL-17 and IL-22 (CD4^+^IL-17^+^IL-22^+^) (Ghoreschi et al., 2010) was drastically increased in *C. rodentium*-infected *Gclc^fl/fl^* mice (Figure 2A, 2D), as was the frequency of “pathogenic” Th17 cells producing IL-17 and IFN-γ (CD4^+^IFN-γ^+^IL-17^+^) (Krausgruber et al., 2016) (Figure S2C). In contrast, the frequency of LP Th1 cells producing IFN-γ but not IL-17 (CD4^+^IFN-γ^+^IL-17^-^) was not increased in infected *Gclc^fl/fl^* mice (Figure 2E). Importantly, IL-17^+^IL-22^+^ pathogenic Th17 cells showed a ∼40-fold increase in absolute cell numbers in response to infection, compared to a ∼4 fold rise in numbers of conventional Th17 cells, Th22 cells, and Th1 cells and a ∼14 fold increase in numbers of IL-17^+^IFN-γ^+^ pathogenic Th17 cells (Figure 2F, S2C). Thus, the response to *C. rodentium* infection in the LP is mediated mainly by pathogenic IL-17^+^IL-22^+^ Th17 cells.

**Figure 2:**
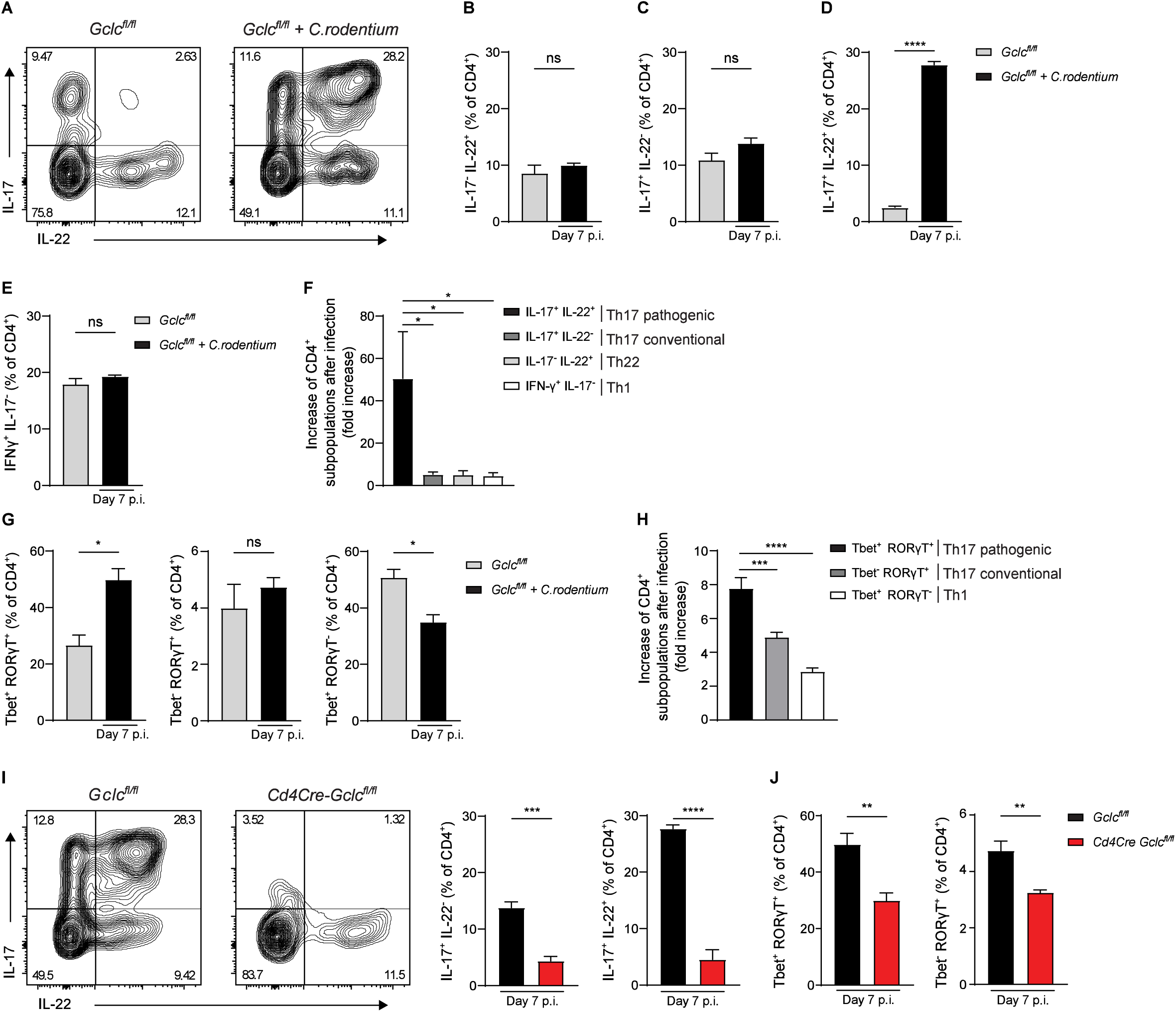
*Gclc* is essential for Th17 cell responses during *C. rodentium* infection. (A) Intracellular FCA of IL-17 and IL-22 in CD4^+^ cells of the colonic LP, which was isolated from uninfected and *C. rodentium*-infected *Gclc^fl/fl^* mice at day 7 p.i. LP cells were stimulated with PMA/calcium ionophore/Brefeldin A for 5 hr before staining. Plots are representative of 4 mice/genotype. (B-E) Frequencies of IL-17^-^IL-22^+^ Th22 cells (B), IL-17^+^IL-22^-^ conventional Th17 cells (C), IL-17^+^IL-22^+^ pathogenic Th17 cells (D), and IFN-γ^+^IL-17^-^ Th1 cells (E) among the colonic LP CD4^+^ cells in (A). Data are mean±SEM (n=4); 3 trials. (F) Quantification of FCA of the indicated CD4^+^ T cell subsets among colonic LP cells from the mice in (A). Relative fold increase represents the ratio of cell numbers in infected/uninfected mice. Data are mean±SEM (n=4); 3 trials. (G) Quantification of intracellular FCA of Tbet and RORγT in CD4^+^ cells of the colonic LP, which was isolated from uninfected and *C. rodentium*-infected *Gclc^fl/fl^* mice at day 7 p.i. Data are frequencies of Tbet^+^RORγT^+^ (left), Tbet^-^RORγT^+^ (middle) and Tbet^+^RORγT^-^ (right) cells among colonic LP CD4^+^ cells. Data are mean±SEM (n=3-6); 2 trials. (H) Quantification of FCA of the indicated CD4^+^ T cell subsets among colonic LP cells from the mice in (G). Relative fold increase represents the ratio of cell numbers in infected/uninfected mice. Data are mean±SEM (n=3-6); 2 trials. (I) Left: Intracellular FCA of IL-17 and IL-22 in CD4^+^ cells of the colonic LP, which was isolated from *C. rodentium*-infected *Gclc^fl/fl^* and *Cd4Cre Gclc^fl/fl^* mice at day 7 p.i. Middle: Frequencies of IL-17^+^IL-22^-^ conventional Th17 cells from left panel results. Right: Frequencies of IL-17^+^IL-22^+^ pathogenic Th17 cells from left panel results. Data are mean±SEM (n=4); 3 trials. (J) Quantification of intracellular FCA of Tbet and RORγT in CD4^+^ cells of the colonic LP, which was isolated from *C. rodentium*-infected *Gclc^fl/fl^* and *Cd4Cre Gclc^fl/fl^* mice at day 7 p.i. Data are frequencies of Tbet^+^RORγT^+^ (left) and Tbet^-^RORγT^+^ (right) cells among colonic LP CD4^+^ cells. Data are mean±SEM (n= 6); 2 trials.

Retinoid orphan receptor RORγT is the master transcription factor (TF) driving Th17 cell differentiation and IL-17 expression, while Tbet is the TF responsible for Th1 cell differentiation and IFN-γ production (Ivanov et al., 2006, Szabo et al., 2000). A bacteria-driven colitis is often associated with pathogenic Th17 cells, which co-express RORγT and Tbet (Krausgruber et al., 2016, Ghoreschi et al., 2010). In colonic LP of *C. rodentium*-infected *Gclc^fl/fl^* mice, the frequency of Tbet^+^RORγT^+^ pathogenic Th17 cells rose sharply, whereas no increase was observed in the frequencies of Tbet^-^RORγT^+^ conventional Th17 cells or Tbet^+^RORγT^-^ Th1 cells (Figure 2G). Total numbers of Tbet^+^RORγT^+^ pathogenic Th17 cells also increased markedly after infection (Figure 2H). These results confirm the previous finding that *C. rodentium* elicits stronger Th17 than Th1 responses (Mangan et al., 2006) and show that specifically the pathogenic Th17 subset, producing both IL-17 and IL-22, is dominant in the T cell response against this pathogen.

### GSH is crucial for the IL-17^+^IL-22^+^ Th17 response during *C. rodentium* infection

*Cd4Cre Gclc^fl/fl^* mice show normal thymocyte development (Mak et al., 2017). Frequencies and absolute numbers of activated/memory T cells, as well as pathogenic and conventional Th17 cells, are comparable in the LP of uninfected *Cd4Cre Gclc^fl/fl^* mice and controls (Figure S2D-F). Thus, *Gclc* is not necessary to maintain these cell populations in mice at steady-state. However, analysis of CD4^+^ T cell subsets from colonic LP of *C. rodentium*-infected *Cd4Cre Gclc^fl/fl^* and *Gclc^fl/fl^* mice on day 7 p.i. showed striking reductions in the frequencies of conventional (IL-17^+^IL-22^-^) and pathogenic (IL-17^+^IL-22^+^ and IFN-γ^+^IL-17^+^) Th17 cells in the mutants (Figure 2I, S2G). The frequency of Th22 cells (IL-17^-^IL-22^+^) was unchanged in infected mutants, and that of Th1 cells (IFN-γ^+^IL-17^-^) was only slightly reduced (Figure S2H, S2I). While absolute numbers of all subsets were decreased in infected *Cd4Cre Gclc^fl/fl^* mice compared to controls, the most prominent effect was on IL-17^+^IL-22^+^ pathogenic Th17 cells (Figure S2J). Frequencies and absolute numbers of Tbet^+^RORγT^+^ pathogenic and Tbet^-^RORγT^+^ conventional Th17 cells were also decreased in LP of infected *Cd4Cre Gclc^fl/fl^* mice (Figure 2J, S2K). No decrease in frequency or absolute number of Tbet^+^RORγT^-^ Th1 cells occurred in infected mutant mice (Figure S2L).

Thus, absence of *Gclc* profoundly affects the IL-17^+^IL-22^+^ pathogenic Th17 cells that dominate responses to *C. rodentium* infection, indicating that *Gclc* is critical for mounting Th17 cell responses to GI infections.

### *Gclc* ablation does not impair the innate lymphoid cell responses to *C. rodentium*

Although IL-17 and IL-22 are signature Th17 cytokines, these mediators are also co-produced by innate lymphoid cells-type 3 (ILC3s) (Satoh-Takayama, 2015). While most ILC3s do not express CD4, a subset called lymphoid tissue inducer (LTi) cells does express CD4 and so would be targeted by *Cd4Cre*-driven *Gclc* deletion in our mouse model. CD4^+^ LTi cells are a critical source of IL-22 in the gut during the early stages of *C. rodentium* infection, when bacteria have just colonized the caecum and innate immunity is dominant (Geddes et al., 2011, Sonnenberg et al., 2011). To determine if loss of LTi function contributed to the phenotype of our infected *Cd4Cre Gclc^fl/fl^* mice during early infection, we analyzed caecal LP CD4^+^ LTi cells (Lineage^-^CD3^-^ Nkp46^-^CD4^+^) from infected *Cd4Cre Gclc^fl/fl^* and *Gclc^fl/fl^* mice at day 4 p.i. by flow cytometry. Neither the frequency nor absolute number of LTi cells nor their cytokine production was affected by *Gclc* ablation (Figure S3A, S3B). The same results were obtained for colonic LP LTi cells at day 4 p.i. (Figure S3C, S3D). When we examined both T cell (CD3^+^CD4^+^) and LTi cell populations at day 7 p.i., when bacteria have reached the colon, the frequencies and absolute numbers of total LTi cells as well as IL-17^+^IL-22^-^, IL-17^-^IL-22^+^ and IL-17^+^IL-22^+^ LTi cells were identical in infected mutant and control mice (Figure 3A-C). In contrast, and consistent with our earlier findings, the frequency and absolute number of IL-17^+^IL-22^+^ Th17 cells in colonic LP were drastically reduced in infected mutants compared to controls (Figure S3E). *Gclc* ablation in colonic LP LTi cells did not affect their intracellular ROS levels, whereas *Gclc*-deficient colonic LP T cells showed highly increased intracellular ROS (Figure 3D, S3F). Thus, contrary to the abrogated IL-22 production seen in Th17 cells, *Gclc* loss does not result in oxidative stress in LP LTi cells and therefore doesn’t interfere with their IL-17 and IL-22 production in *C. rodentium*-infected mice.

**Figure 3:**
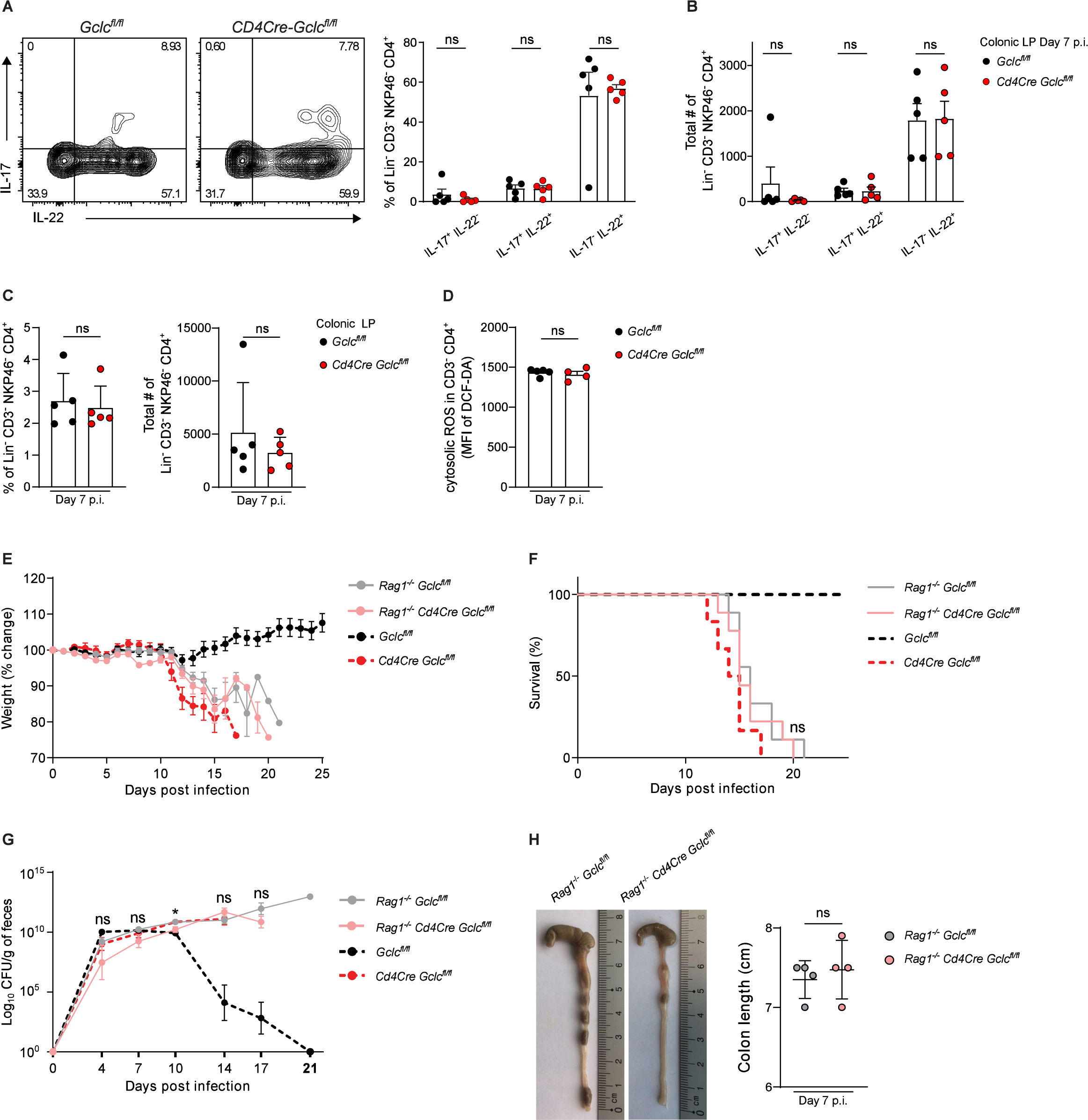
Absence of *Gclc* in LTi cells does not impair cytokine production or immunity to *C. rodentium*. (A) Left: Intracellular FCA of IL-17 and IL-22 in colonic LP LTi cells of *C. rodentium*-infected *Gclc^fl/fl^* and *Cd4Cre Gclc^fl/fl^* mice at day 7 p.i. LP cells were stimulated with PMA/calcium ionophore/Brefeldin A for 5 hr before staining. Right: Frequencies of the indicated subpopulations from left panel results. Data are mean±SEM (n=5); 3 trials. (B) Total cell numbers of the indicated subpopulations from the data in (A). Data are mean±SEM (n=5); 3 trials. (C) Frequencies (left) and total numbers (right) as determined by FCA of LTi cells among colonic LP cells isolated from *C. rodentium*-infected *Gclc^fl/fl^* and *Cd4Cre Gclc^fl/fl^* mice at day 7 p.i. Data are mean±SEM (n=5); 3 trials. (D) Quantification of FCA of intracellular ROS in CD3^-^CD4^+^ LTi cells of the colonic LP, which was isolated from *C. rodentium*-infected *Gclc^fl/fl^* and *Cd4Cre Gclc^fl/fl^* mice at day 7 p.i. and subjected to DCF-DA staining. Data are mean±SEM (n=4-5). (E) Change in whole body weight of *Rag1^-/-^Gclc^fl/fl^*, *Rag1^-/-^Cd4Cre Gclc^fl/fl^, Gclc^fl/fl^* and *Cd4Cre Gclc^fl/fl^* mice during *C. rodentium* infection. Data are mean±SEM (n=6-9); 3 trials. (F) Survival of *Rag1^-/-^Gclc^fl/fl^* (n=9), *Rag1^-/-^Cd4Cre Gclc^fl/fl^* (n=9)*, Gclc^fl/fl^* (n=6) and *Cd4Cre Gclc^fl/fl^* (n=6) mice during *C. rodentium* infection. Data are representative of 3 trials. (G) Left: CFU of *C. rodentium* in feces of *Rag1^-/-^Gclc^fl/fl^*, *Rag1^-/-^Cd4Cre Gclc^fl/fl^, Gclc^fl/fl^* and *Cd4Cre Gclc^fl/fl^* mice at the indicated time points after infection on day 0. Right: Statistical analysis of left panel results at day 14 p.i. Data are mean±SEM (n=6-9); 3 trials. (H) Left: Macroscopic views and lengths of colons isolated from *C. rodentium*-infected *Rag1^-/-^ Gclc^fl/fl^* and *Rag1^-/-^Cd4Cre Gclc^fl/fl^* mice at day 7 p.i. Right: Quantification of colon lengths from left panel results. Data are mean±SEM (n=4).

To investigate if an effect of *Gclc* ablation in LTi cells might be masked by an adaptive immune response, we crossed *Cd4Cre Gclc^fl/f^* mice with *Rag1^-/-^* mice to generate a *Rag1^-/-^Cd4Cre Gclc^fl/fl^* strain lacking the adaptive immune system (Mombaerts et al., 1992). Once again, the frequencies and absolute numbers of IL-17^+^IL-22^-^, IL-17^-^IL-22^+^ and IL-17^+^IL-22^+^ LTi cells in colonic LP, as well as their intracellular ROS levels, were identical in *C. rodentium*-infected *Rag1^-/-^Cd4Cre Gclc^fl/fl^* and control *Rag1^-/-^Gclc^fl/fl^* mice (Figure S3G, S3H). When we compared the course of disease in infected *Rag1^-/-^Cd4Cre Gclc^fl/fl^, Rag1^-/-^Gclc^fl/fl^*, *Cd4Cre Gclc^fl/fl^* and *Gclc^fl/fl^* mice, we observed that both *Rag1^-/-^Cd4Cre Gclc^fl/fl^* and *Rag1^-/-^Gclc^fl/fl^* mice exhibited severe weight loss and succumbed to the infection by day 15-20 p.i. (Figure 3E, 3F), consistent with the documented importance of the adaptive response in controlling the later stages of *C. rodentium* infection (Vallance et al., 2002, Bry and Brenner, 2004, Sonnenberg et al., 2011). However, no differences in survival, weight loss, or fecal bacterial load were observed between *Rag1^-/-^Cd4Cre Gclc^fl/fl^* and *Rag1^-/-^Gclc^fl/fl^* mice at day 4 p.i., when protection is primarily mediated by innate immune cells (Figure 3E–3G). In addition, colon appearance and length were comparable between infected *Rag1^-/-^Cd4Cre Gclc^fl/fl^* and *Rag1^-/-^Gclc^fl/fl^* mice at day 7 p.i. (Figure 3H). Thus, the innate response is intact in *Cd4Cre Gclc^fl/fl^* mice and their vulnerability to *C. rodentium* infection is due to their lack of T cell function.

### GSH-regulated IL-22 is essential for clearance of *C. rodentium*

IL-17, IL-22, IFN-γ and TNF-α all contribute to controlling the spread of *C. rodentium* within the body and limiting intestinal epithelial damage (Mangan et al., 2006, Ishigame et al., 2009, Basu et al., 2012, Zheng et al., 2008, Simmons et al., 2002, Shiomi et al., 2010, Gonçalves et al., 2001). However, while mice lacking IL-17, IFN-γ or TNF-α all show increased histological disease scores and high fecal bacterial titers, only mice lacking IL-22 are actually unable to clear *C. rodentium* and succumb to the infection (Basu et al., 2012, Zheng et al., 2008, Ota et al., 2011). Because IL-17^+^IL-22^+^ Th17 cells were greatly reduced in the colonic LP of our infected *Cd4Cre Gclc^fl/fl^* mice, we hypothesized that loss of Th17 cell-derived IL-22 caused their high mortality. We therefore treated infected *Cd4Cre Gclc^fl/fl^* and *Gclc^fl/fl^* mice with either a recombinant IL-22-Fc fusion protein or a control-Fc fragment and monitored mouse weight and survival. Infected mutant mice treated with control-Fc showed severe weight loss and 100% mortality, whereas recombinant IL-22-Fc treatment completely prevented weight loss and fully restored survival (Figure 4A, 4B). Strikingly, IL-22-Fc treatment also blocked the intestinal ulceration and crypt loss observed in control-Fc-treated *Cd4Cre Gclc^fl/fl^* mice (Figure 4C). We therefore speculated that, because the *Gclc-*deficient T cells in the LP of infected mutants produced very little IL-22, systemic effects were triggered that led to a fatal outcome.

**Figure 4:**
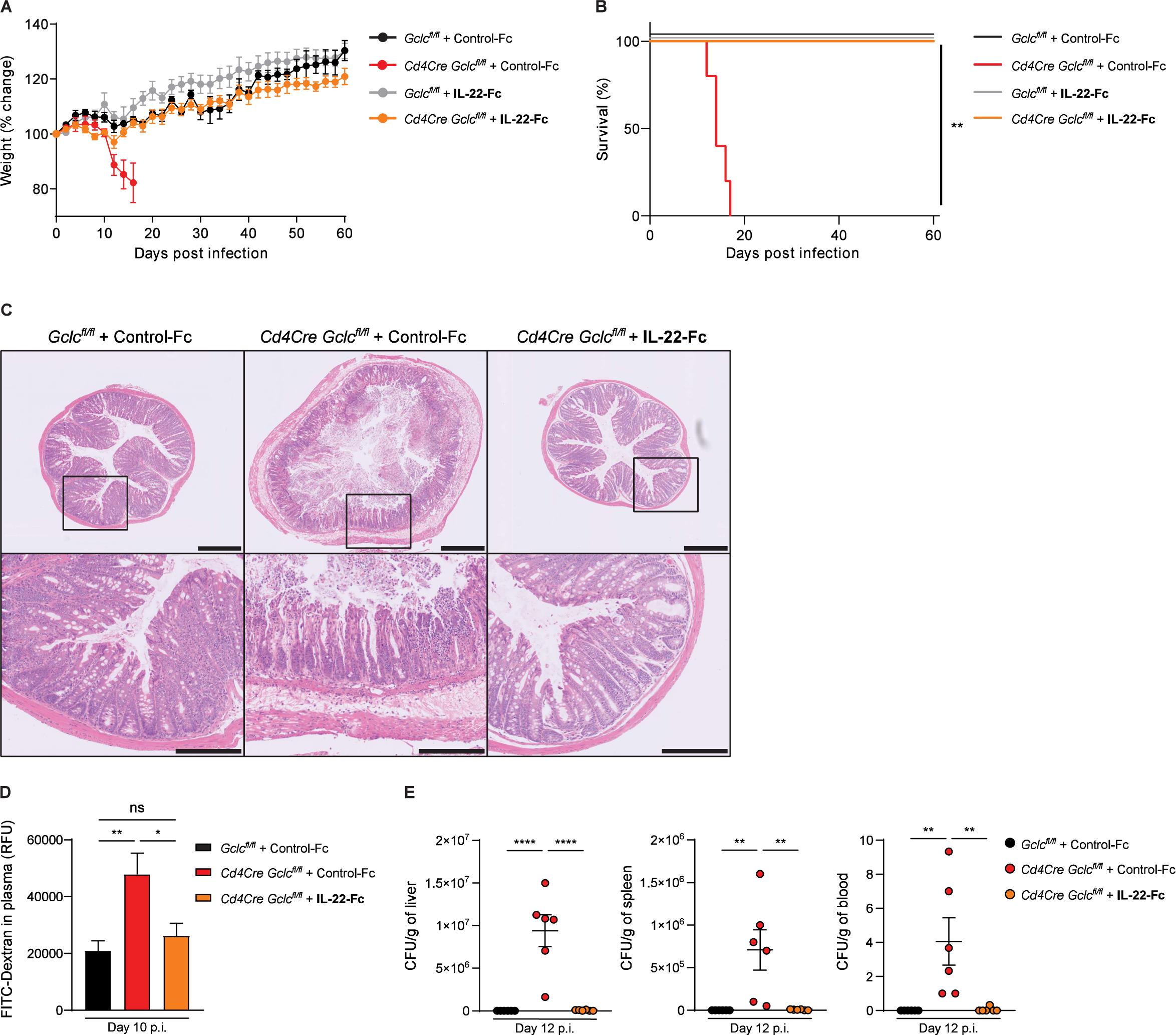
IL-22 reconstitution rescues mice with T cell-specific GSH deficiency from lethal *C. rodentium* infection and prevents increased intestinal permeability. (A) Change in whole body weight of *C. rodentium*-infected *Gclc^fl/fl^* and *Cd4Cre Gclc^fl/fl^* mice treated i.v. with recombinant IL-22-Fc or control-Fc at days −1, 3, 6, 9, 12 and 15 p.i. Mice were infected on day 0. Data are mean±SEM (n=4-5); 2 trials. (B) Survival of the mice in (A). Data are representative of 2 trials. (C) Top: H&E-stained sections of distal colon of *C. rodentium*-infected *Gclc^fl/fl^* and *Cd4Cre Gclc^fl/fl^* mice that were treated i.v. with IL-22-Fc or control-Fc as in (A, B) and examined at day 12 p.i. Data are representative of 3 mice/genotype. Scale bars, 500µm. Bottom: Higher magnification views of the boxed areas in the top panels. Scale bars, 200µm. (D) Quantification of plasma FITC-Dextran levels in mice of the indicated genotypes that were infected with *C. rodentium* and treated i.v. with IL-22-Fc or control-Fc at days −1, 3, 6, 9, p.i. At day 10 p.i., mice were gavaged with FITC-Dextran. FITC-Dextran was measured in plasma 4 hr later. Data are mean±SEM (n=6). (E) CFU of *C. rodentium* in liver (left), spleen (middle) and blood (right) of *C. rodentium*-infected *Gclc^fl/fl^* and *Cd4Cre Gclc^fl/fl^* mice treated i.v. with IL-22-Fc or control-Fc at days −1, 3, 6, 9 p.i. Organs and blood were collected at day 12 p.i. Data are mean±SEM (n=6).

### *Gclc*-dependent IL-22 regulates intestinal permeability during gut infection

IL-22 is involved in many pathways that influence anti-bacterial immunity, including those affecting pathogen virulence (Sakamoto et al., 2017, Sanchez et al., 2018). To investigate whether *C. rodentium* virulence was altered in infected *Cd4Cre Gclc^fl/fl^* mice, we determined mRNA levels of the bacterial virulence factors EspA, EspG, EspF, EspI, Tir and Map in colonic tissues of *Cd4Cre Gclc^fl/fl^* and *Gclc^fl/fl^* mice at day 7 or 10 p.i. However, we found no differences between mutant and control mice (Figure S4A) indicating that IL-22 lacking in *Cd4Cre Gclc^fl/fl^* T cells does not impact bacterial virulence.

IL-22 also regulates IEC production of AMPs such as Reg and β-defensins as well as lipocalin-2 (Liang et al., 2006, Zheng et al., 2008, Raffatellu et al., 2009, Dixon et al., 2016). We therefore measured mRNA levels of RegIIIβ, RegIIIγ, β-defensin-2 and lipocalin-2 in caecum and distal colon of infected *Cd4Cre Gclc^fl/fl^* and *Gclc^fl/fl^* mice. Again, no major differences were detected between mutant and control mice (Figure S4B, S4C). Moreover, lipocalin-2 levels in feces were comparable between mutant and control mice throughout the infection (Figure S4D).

IL-22 has also been linked to intestinal barrier integrity and permeability (Keir et al., 2020, Rendon et al., 2013). To measure intestinal permeability, we orally administered fluorescein isothiocyanate (FITC)-Dextran to *Cd4Cre Gclc^fl/fl^* and *Gclc^fl/fl^* mice on day 10 p.i. and measured FITC-Dextran accumulation in blood plasma. Permeation of FITC-Dextran from the intestinal lumen into the plasma was greatly enhanced in infected mutant mice compared to controls (Figure S4E), but comparable in uninfected mutant and control mice (Figure S4F). This increase in intestinal permeability was prevented by IL-22-Fc treatment of *Cd4Cre Gclc^fl/fl^* mice, indicating restoration of an intact intestinal layer (Figure 4D). By day 12 p.i., we detected high bacterial titers in the blood, spleen and liver of infected *Cd4Cre Gclc^fl/fl^* mice, whereas infected control mice showed a minimal bacterial burden (Figure S4G, 4E). Again, IL-22-Fc treatment of infected mutant mice reduced their bacterial loads back to control levels (Figure 4E). These data point to a mechanism of *Gclc*-dependent regulation of T cell-intrinsic IL-22 production that protects intestinal integrity and so prevents bacterial spread from the intestinal lumen to the periphery.

### *Gclc*-regulated IL-22 bolsters intestinal tight junctions and mucus layer in *C. rodentium*-infected mice

IL-22 contributes to intestinal integrity by regulating the expression and intracellular location of tight junction proteins such as the claudins, which maintain the epithelial barrier (Kim et al., 2012, Tsai et al., 2017). Intact tight junctions are crucial for protection against infection by attaching/effacing pathogens such as *C. rodentium*, which specifically target tight junction structural proteins (Guttman et al., 2006, Garber et al., 2018, Xia et al., 2019). To investigate whether *Gclc* ablation in T cells decreases claudin proteins, we immunoblotted distal colon sections of *Cd4Cre Gclc^fl/fl^* and *Gclc^fl/fl^* mice at day 12 p.i. Indeed, colons from infected mutants showed lower claudin-2, claudin-3 and claudin-15 protein levels compared to colons from infected controls (Figure 5A). In line, immunofluorescence microscopy of cross-sections of colonic crypts showed abnormally diffused claudin-2 localization in infected *Cd4Cre Gclc^fl/fl^* mice (Figure 5B). Proper localization of claudin-2 in colonic crypts of infected *Cd4Cre Gclc^fl/fl^* mice, as well as normal lumen diameters, were entirely restored after IL-22-Fc treatment (Figure 5B).

**Figure 5:**
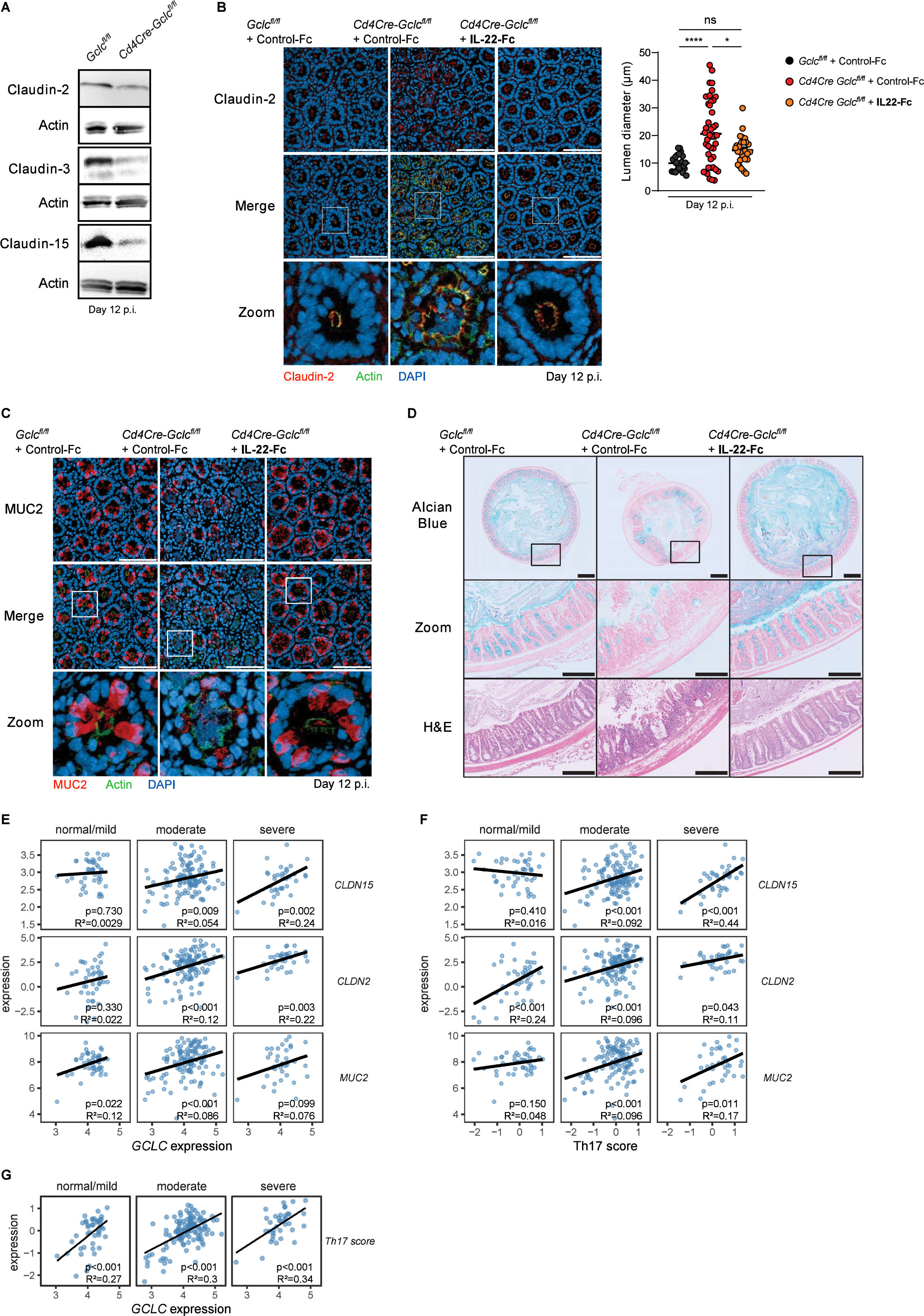
*Gclc* deficiency is linked to defects in intestinal tight junctions and mucus in *C. rodentium*-infected mice and IBD patients. (A) Representative immunoblots to detect the indicated claudin proteins in distal colon segments of *C. rodentium*-infected *Gclc^fl/fl^* and *Cd4Cre Gclc^fl/fl^* mice at day 12 p.i. Actin, loading control. Data are representative of 3 mice/genotype. (B) Left: Top and middle: Immunofluorescence microscopy of cross-sections of colonic crypts from distal colon segments that were collected at day 12 p.i. from *C. rodentium*-infected *Gclc^fl/fl^* and *Cd4Cre Gclc^fl/fl^* mice treated i.v. with IL-22-Fc or control-Fc at days −1, 3, 6, 9, p.i. Sections were stained to detect claudin-2 (red), F-actin (green) and DNA (blue). Data are representative of 6 mice/genotype. Scale bars, 100µm. Bottom: “Zoom” panels are higher magnification views of boxed areas in the middle panels. Right: Quantification of lumen diameters of 24-43 crypts measured per mouse in the left panel. Data are representative of 6 mice/genotype. (C) Top and middle: Immunofluorescence microscopy of cross-sections of colonic crypts from distal colon segments in (B) were stained to detect MUC2 (red), F-actin (green) and DNA (blue). Data are representative of 6 mice/genotype. Scale bars, 100µm. Bottom: “Zoom” panels are higher magnification views of boxed areas in the middle panel. (D) Top: Alcian blue-stained cross-sections of distal colon segments that were collected at day 12 p.i. from *C. rodentium*-infected *Gclc^fl/fl^* and *Cd4Cre Gclc^fl/fl^* mice treated i.v. with IL-22-Fc or control-Fc at days −1, 3, 6, 9, p.i. Data are representative of 3 mice/genotype. Scale bars, 500µm. Middle: “Zoom” panels are higher magnification views of the boxed areas in the top panels. Scale bars, 200µm. Bottom: H&E staining of the sections in the middle panels. Scale bars, 200µm. (E, F) Bioinformatics analyses of RNA sequencing data of rectal biopsies from ulcerative colitis (UC) patients. The provided MAYO scores were used to define the disease severity as normal/mild, moderate and severe disease (Teixeira et al., 2015). Significance and r^2^ values are indicated in the lower right corner of each frame. (E) Correlation of *GCLC* expression with the epithelial integrity markers claudin-15 (*CLDN15*) (top), claudin-2 (*CLDN2*) (middle), and mucin 2 (*MUC2*) (bottom). (F) Correlation of the Th17 score with the epithelial integrity markers claudin-15 (*CLDN15*) (top), *CLDN2* (middle), and *MUC2* (bottom). The Th17 score was defined as the mean of scaled log2 gene expression values composing the signature (*IL-17A, IL-17F, IFN-γ, CD3E, CD4, STAT3, STAT5a, STAT1, STAT4, STAT6, AHR, RORC, RORA and TBX21*). (G) Bioinformatics analysis of RNA sequencing data of UC rectal biopsies as in (E,F) showing correlation of *GCLC* expression with Th17 score.

Another key function of IL-22 associated with intestinal pathogen exclusion is promotion of mucin production by intestinal goblet cells (Sugimoto et al., 2008). The mucin-containing mucus layer that protects the intestinal epithelium from enteric contents is critical for mouse survival of *C. rodentium* infection (Sugimoto et al., 2008, Bergstrom et al., 2010, Desai et al., 2016). The mucin protein MUC2 is the main component of the intestinal mucus layer and most abundant in the colon (Johansson et al., 2008). We performed immunofluorescence microscopy of cross-sections of intestinal crypts and observed a great reduction in MUC2 in

IECs of infected *Cd4Cre Gclc^fl/fl^* mice on day 12 p.i. (Figure 5C). Again, normal MUC2 levels were completely restored by IL-22-Fc treatment (Figure 5C). Alcian blue staining of transverse sections of distal colons revealed a drastic decrease in total luminal mucus and the adherent mucus layer in colons of infected *Cd4Cre Gclc^fl/fl^* mice compared to controls (Figure 5D). Moreover, the amount of mucin stored inside colonic IECs was lower in control-Fc-treated *Cd4Cre Gclc^fl/fl^* mice; especially in sites of ulceration (Figure 5D). Once again, mucus erosion was prevented in infected mutant mice by IL-22-Fc injection (Figure 5D). These data indicate the existence of a crucial axis between T cell-intrinsic GSH synthesis and IL-22 production that is indispensable for the protective function of the intestinal epithelial barrier.

### *GCLC* expression in IBD patients correlates positively with expression of genes related to gut integrity

Our results above prompted us to investigate whether *GCLC* expression could be linked to intestinal integrity in IBD patients. To this extend, we analyzed publicly available RNA sequencing dataset (GSE109142) of rectal biopsies from patients with ulcerative colitis (UC), a subtype of IBD (Haberman et al., 2019). MAYO scores (ranging from 0-12) were used to define the disease severity as normal/mild (score <6), moderate (score <11) and severe (score ≥11) (Teixeira et al., 2015). We then classified the rectal tissues of these patients as having low or high *GCLC* expression (Figure S5A). In patinets with moderate or severe disease, we identified a positive correlation between *GCLC* expression and that of *CLDN15* (claudin-15) and *CLND2* (claudin-2) (Figure 5E, S5B). *MUC2* expression correlated positively with *GCLC* in patients with mild or moderate disease; patients with severe pathology likewise displayed a positive slope but the correlation was not significant (Figure 5E, S5B). These data from IBD patients are in line with our mouse results and further support our hypothesis that *Gclc* expression plays a key role in preventing gut pathology.

In mouse models, IL-17 and IL-22, which can be largely produced by Th17 cell, have been linked to the regulation of claudins and mucins production (Lee et al., 2015, Tsai et al., 2017, Kim et al., 2012, Sugimoto et al., 2008). To investigate this relationship in the human setting, we defined a Th17 cell gene expression signature that included *IL-17A, IL-17F, IFN-γ, CD3E, CD4, STAT3, STAT5a, STAT1, STAT4, STAT6, AHR, RORC, RORA and TBX21*. We observed a strong positive correlation between the Th17 gene signature score and *CLDN15*, *CLND2* and *MUC2* levels in patients with moderate or severe pathology (Figure 5F, S5B). This score also correlated positively with *IL-22* expression, indicating that most of the Th17 cells analyzed expressed both IL-17 and IL-22 (Figure S5C). Thus, these results suggest a link between the expression of Th17-related genes and intestinal epithelial integrity in IBD patients. Notably, although *GCLC* expression did not correlate with disease severity, the expression of glutathione synthase (*GSS*), which catalyzes the second step of GSH synthesis, was significantly reduced in the gut of UC patients with severe pathology (Figure S5D). Moreover, there was a strong positive correlation between *GCLC* expression and the Th17 gene signature score in IBD patients (Figure 5G). These findings are in line with our murine data and indicate a strong association between *GCLC*, Th17 cell markers, and genes related to gut integrity in the intestine of IBD patients.

### T cell-intrinsic IL-22 is sufficient to restore survival of *C. rodentium*-infected *Cd4^Cre^-Gclc^fl/fl^* mice

We showed that systemic injection of recombinant IL-22-Fc prevented *C. rodentium*-induced intestinal damage and mortality in *Cd4Cre Gclc^fl/fl^* mice. To confirm that it is indeed GSH-controlled IL-22 production by T cells that is the protective mechanism, we generated a novel Cre recombinase-inducible IL22 transgenic mouse model (*IL22^ind^* mice) by genetic targeting of embryonic stem cells (Figure S6A, S6B). Animal derived from these cells harbor a transgene in the *Rosa26* locus that contains cDNAs for IL22 and the EGFP reporter preceded by a loxP-flanked STOP cassette and the CAG promoter (Figure S6A, S6B). After Cre-dependent removal of the STOP sequence, IL22 and EGFP can be constitutively expressed. To reconstitute IL-17 production as well, we took advantage of an IL-17 transgenic mouse model where IL-17 and EGFP expression could be controlled in a similar way (Haak et al., 2009). After Cre-dependent removal of the STOP sequence, IL-22 or IL-17A and EGFP can be constitutively expressed. To generate mice with T cell-specific IL-22 or IL-17A expression, we crossed *IL-22^ind^* and *IL-17A^ind^* mice with *Cd4Cre* mice. EGFP and IL-22 were expressed in T cells of *Cd4Cre IL-22^ind/+^* mice, but not in *IL-22^ind/+^* controls (Figure S6C, S6D). Similarly, EGFP and IL-17A were expressed in T cells of *Cd4Cre IL-17A^ind/+^* mice, but not in *IL-17A^ind/+^* controls (Figure S6F). We then crossed *IL-22^ind^* and *IL-17A^ind^* mice with *Cd4Cre Gclc^fl/fl^* mice to generate *Cd4Cre Gclc^fl/fl^ IL-22^ind/+^* and *Cd4Cre Gclc^fl/fl^ IL-17A^ind/+^* mice, which undergo T cell-specific *Gclc* deletion in parallel with induction of EGFP and IL-22 or IL-17A, respectively. This approach uncouples IL-22 and IL-17A expression from its regulation by *Gclc* in endogenous CD4^+^ cells (Figure S6E, S6G).

To investigate whether reinstating IL-22 production specifically in T cells of *Cd4Cre Gclc^fl/fl^* mice was sufficient to prevent infection-induced mortality, we infected *Cd4Cre Gclc^fl/fl^IL-22^ind/+^* and *Cd4Cre Gclc^fl/fl^* mice, along with *Gclc^fl/fl^* and *Gclc^fl/fl^IL-22^ind/+^* controls, with *C. rodentium*. Indeed, transgenetic expression of IL-22 expression in mutant T cells (*Cd4Cre Gclc^fl/fl^IL-22^ind/+^*) rescued these mice from the lethal infection outcome to a large extent and mice were protected from weight loss (Figure 6A, S6H). When not completely protected from infection-induced lethality *Cd4Cre Gclc^fl/fl^IL-22^ind/+^* showed a prolonged survival compared to infected *Cd4Cre-Gclc^fl/fl^* mice (Figure 6A). *Cd4Cre-Gclc^fl/fl^IL-22^ind/+^* mice were able to clear *C. rodentium* infection by around day 60 p.i. (Figure S6I). In contrast to IL-22, reinstating IL-17 expression in mutant T cells did not increase survival, nor could it limit weight loss (Figure 6B, S6J, K). Importantly, T cell-specific expression of IL-22, in mutant T cells prevented bacterial spreading to spleen and liver, which was in stark contrast to the transgenetic expression of IL-17 in *Gclc-* deficient (Figure 6C). These perplexing results indicate that it is the GSH-controlled T cell-derived IL-22 and not the T cell-derived IL-17, which is sufficient to support full recovery of these mutants. However, bacterial clearance in *Cd4Cre Gclc^fl/fl^IL-22^ind/+^* took more than twice as long as in controls. This emphasizes the existence of additional factors that are under control of *Gclc*, which contribute to optimal bacterial clearance. Nevertheless, our data indicate that GSH-regulated IL-22 production by T cells appears to be the major factor protecting the gut from detrimental bacterial infections.

**Figure 6:**
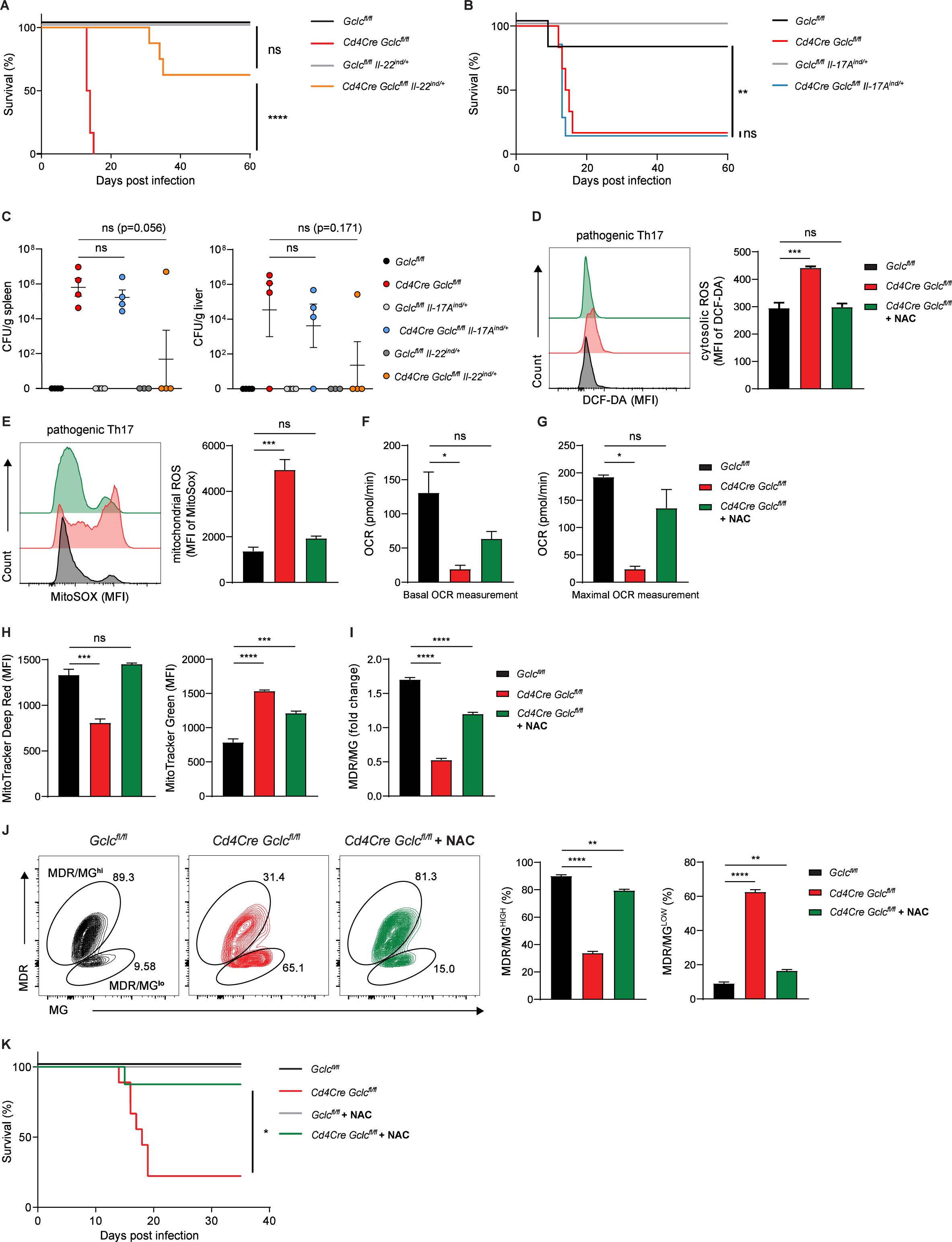
T-cell intrinsic IL-22 but not IL-17 expression rescues *Gclc*-dependent susceptibility to *C.rodentium*. (A) Survival of *Gclc^fl/fl^* (n=5), *Cd4Cre Gclc^fl/fl^* (n=6)*, Gclc^fl/fl^*IL-22^ind/+^ (n=5) and *Cd4Cre Gclc^fl/fl^*IL-22^ind/+^ (n=8) mice infected with *C. rodentium* on day 0 and assayed on the indicated days p.i. Data are mean±SEM (n=5-8). (B) Survival of of *Gclc^fl/fl^* (n=6), *Cd4Cre Gclc^fl/fl^* (n=9)*, Gclc^fl/fl^*IL-17A^ind/+^ (n=6) and *Cd4Cre Gclc^fl/fl^*IL-17A^ind/+^ (n=7) mice infected with *C. rodentium* on day 0 and assayed on the indicated days p.i. Data are mean±SEM (n=6-9). (C) CFU of *C. rodentium* in spleen (left) and liver spleen (right) of *C. rodentium*-infected *Gclc^fl/fl^*, *Cd4Cre Gclc^fl/fl^, Gclc^fl/fl^*IL-22^ind/+^, *Cd4Cre Gclc^fl/fl^*IL-22^ind/+^*, Gclc^fl/fl^*IL-17A^ind/+^ and *Cd4Cre Gclc^fl/fl^*IL-17A^ind/+^ mice (n=4), day 10 p.i. (D-J) Naïve T cells were sorted from spleen and lymph nodes of *Gclc^fl/fl^* and *Cd4Cre Gclc^fl/fl^* mice and induced to differentiate *in vitro* into pathogenic Th17 cells by culture with anti-CD3, anti-CD28, IL-6, IL-1β and IL-23. Cells were treated with 10mM NAC as indicated. (D) Left: FCA of DCF-DA staining to detect cytosolic ROS in *Gclc^fl/fl^* and *Cd4Cre Gclc^fl/fl^ in vitro-* differentiated Th17 cells. Right: Quantification of left panel results. Data are mean±SEM (n=3); 2 trials. (E) Left: FCA of MitoSOX to detect mitochondrial ROS in *Gclc^fl/fl^* and *Cd4Cre Gclc^fl/fl^ in vitro*-differentiated Th17 cells. Right: Quantification of left panel results. Data are mean±SEM (n=3); 2 trials. (F, G) Seahorse quantification of basal OCR (E) and maximal OCR (F) in cultures of *Gclc^fl/fl^* and *Cd4Cre Gclc^fl/fl^ in vitro*-differentiated Th17 cells. Data are mean±SEM (n=3); 2 trials. (H) Quantification of FCA of *Gclc^fl/fl^* and *Cd4Cre Gclc^fl/fl^ in vitro*-differentiated Th17 cells that were treated with (left) MitoTracker Deep Red to assess mitochondrial activity, or (right) MitoTracker Green to assess mitochondrial mass. Data are mean±SEM (n=3); 3 trials. (I) Determination of fold change in the MDR/MG ratio from the data in (B). Data are mean±SEM (n=3); 3 trials. (J) Representative contour plots (left) and frequencies (right) of MDR/MG^hi^ and MDR/MG^lo^ cell subpopulations among *in vitro*-differentiated *Gclc^fl/fl^* and *Cd4Cre Gclc^fl/fl^* Th17 cultures as determined by FCA. Data are mean±SEM (n=3); 3 trials. (K) Survival of *C. rodentium*-infected *Gclc^fl/fl^* and *Cd4Cre Gclc^fl/fl^* mice (n=5-9) that were left untreated or treated with 40mM NAC in drinking water. Data are pooled from 2 independent trials.

### Mitochondrial function is impaired in *Gclc*-deficient Th17 cells

To get more mechanistic insights how GSH regulates IL-22 production in Th17 cells we took advantage of *in vitro* differentiated Th17 cells. Naïve CD4^+^ T cells from *Cd4Cre Gclc^fl/fl^* mice were activated in the presence of IL-6, IL-23 and IL-1β and skewed to Th17 cells. These conditions are known to induce naïve CD4^+^ T cells to differentiate into a pathogenic Th17 cell subset co-producing IL-17 and IL-22 (Budda et al., 2016, Ghoreschi et al., 2010). In contrast, conventional Th17 cells producing mainly IL-17 are induced if naive CD4^+^ T cells are activated in the presence of IL-6 and TGF-β, since TGF-β inhibits IL-22 production (Rutz et al., 2011). In line with our *in vivo* data, we found that *Cd4Cre-Gclc^fl/fl^* CD4^+^ T cells differentiated into pathogenic Th17 cells produced less IL-22 and IL-17 than controls (Figure S7A, B). In addition, mutant Th17 cells showed increased cytosolic and mitochondrial ROS due to a lack of ROS buffering caused by their GSH deficiency (Figure 6D, E). Accordingly, cytosolic and mitochondrial ROS were greatly increased in *Gclc*-deficient LP T cells isolated from mice at 7 days p.i. with *C. rodentium* (Figure S7C, D). The addition of the antioxidant N-acetyl-cysteine (NAC) during *in vitro* differentiation of the mutant cells restored ROS to control levels (Figure 6D, E). The same results were obtained when conventional Th17 cells were generated (Figure S7E), implying that ROS accumulate in *Gclc*-deficient Th17 cells regardless of the differentiation conditions.

Accumulating ROS can impact the mitochondrial dynamics and function (Chakrabarty and Chandel, 2021). To assess the functionality of mitochondria in mutant Th17 cells, we measured the oxygen consumption rate (OCR) in cultures of these cells. Basal and maximal OCR measurements were highly reduced in *Gclc*-deficient pathogenic Th17 cells, but both parameters were partially restored through ROS-scavenging by NAC (Figure 6F, 6G). We then assessed mitochondrial mass (MM) and membrane potential (indicating activity; MMP) in these cells by using MitoTracker Green (MG) and MitoTracker Deep Red (MDR), respectively. Intriguingly, the mutant Th17 cells showed a drastic increase in MM, whereas MMP was dramatically decreased (Figure 6H). The MDR/MG ratio indicates MMP/MM and represents mitochondrial capacity (Pendergrass et al., 2004, Yu et al., 2020b). We found that MDR/MG was decreased in *Gclc*-deficient Th17 cells, suggesting that their mitochondrial capacity is impaired (Figure 6I). A decreased MDR/MG ratio was also observed in *Gclc*-deficient LP T cells isolated at day 7 p.i. (Figure S7F). In addition, most control *Gclc^fl/fl^* Th17 cells constituted a MDR/MG^hi^ population, whereas mutant Th17 cells formed a MDR/MG^lo^ population (Figure 6J), which suggests their mitochondria being depolarized and dysfunctional (Yu et al., 2020b). Mitochondrial polarization and mass in mutant Th17 cells were almost completely restored by NAC (Figure 6H-6J), indicating that mitochondrial activity in these cells depends on ROS control. These results prompted us to investigate whether ROS-scavenging *in vivo* might restore the functionality of mutant Th17 cells. Therefore, we administered NAC in the drinking water of *Cd4Cre-Gclc^fl/fl^* and *Gclc^fl/fl^* mice starting at 7 days before *C. rodentium* infection and continuing throughout the infection. In line with our *in vitro* data, the mortality of NAC-treated infected mutant mice was drastically reduced (Figure 6K).

### *Gclc* regulates the mitochondrial-encoded components of the electron transport chain of Th17 cells

Our data so far indicate that mitochondrial dynamics and activity are affected by the absence of *Gclc* in Th17 cells. To investigate the components of mitochondrial machinery on a larger scale, we conducted bulk RNA sequencing of *in vitro* differentiated pathogenic Th17 cells. Although, mutant Th17 cells showed decreased OXPHOS most of the genes that encoded for subunits of the mitochondrial electron transport chain (ETC) appeared to be significantly upregulated when compared to control cells (Figure 7A). Intriguingly, a closer examination exposed that the ETC genes encoded within mitochondrial DNA were specifically down-regulated in the mutant Th17 cells (Figure 7A). Notably, treating *Gclc-*deficient Th17 cells with NAC restored the expression levels of these mitochondria-encoded ETC components (Figure 7A). This emphasizes that specifically mitochondrial gene expression is negatively impacted by the absence of GSH in Th17 cells.

**Figure 7:**
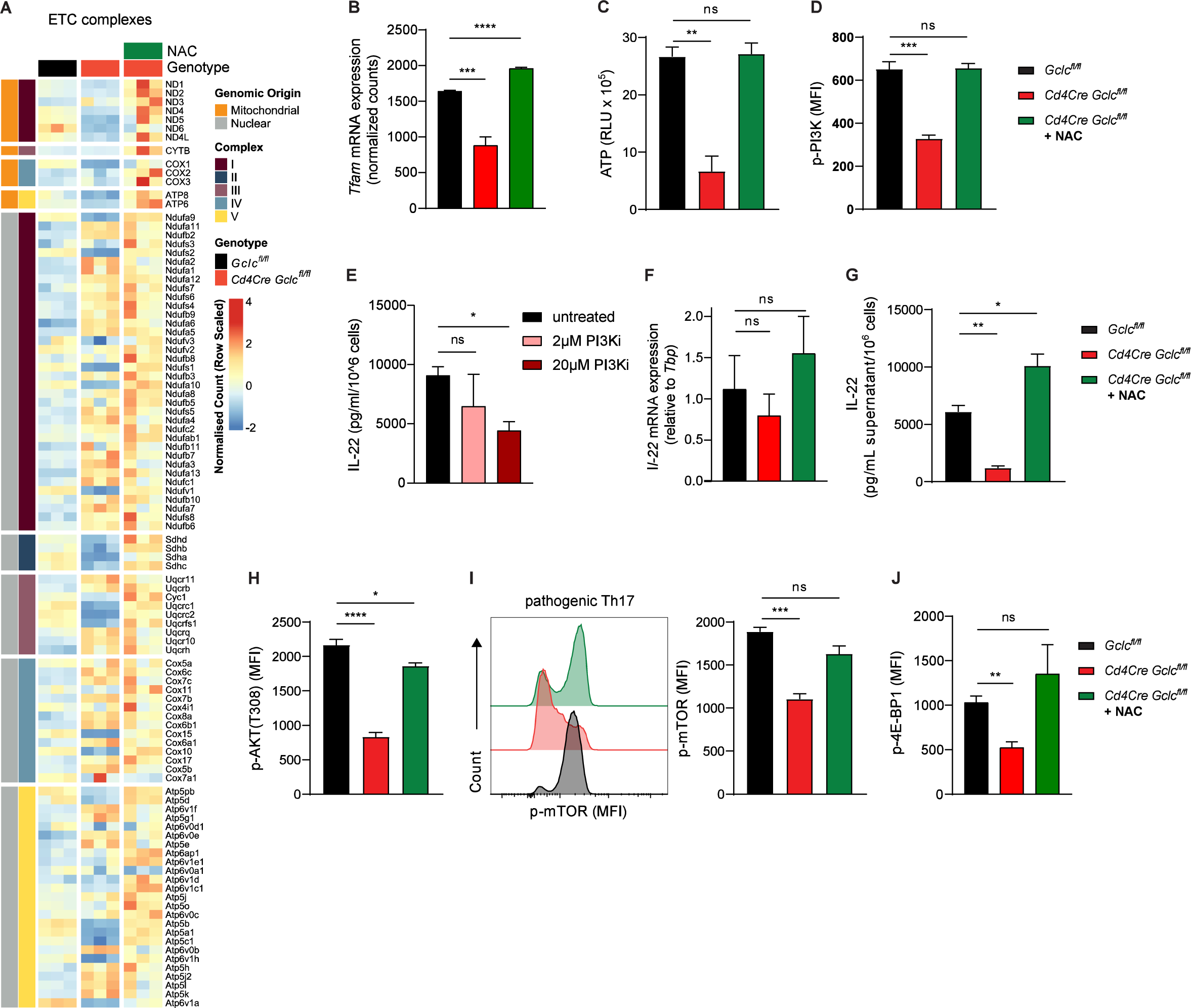
*Gclc* connects mitochondrial gene expression and mitochondrial function with IL-22 protein translation in Th17 cells. (A-J) Naïve T cells were sorted from spleen and lymph nodes of *Gclc^fl/fl^* and *Cd4Cre Gclc^fl/fl^* mice and induced to differentiate *in vitro* into pathogenic Th17 cells by culture with anti-CD3, anti-CD28, IL-6, IL-1β and IL-23. Cells were treated with 10mM NAC as indicated. (A-B) Bulk RNA seq was performed. (A) Heatmap representing GO:0022900 Electron transport Chain. (B) Tfam expression. (n=3). (C) Quantification of total ATP levels in *Gclc^fl/fl^* and *Cd4Cre Gclc^fl/fl^ in vitro*-differentiated Th17 cells as measured in RLU. Data are mean±SEM (n=3); 2 trials. (D) Quantification of FCA of p-PI3K in *in vitro-*differentiated *Gclc^fl/fl^* and *Cd4Cre Gclc^fl/fl^* Th17 cells. Data are mean±SEM (n=3); 2 trials. (E) IL-22 protein concentrations as measured by ELISA in culture supernatants of *in vitro*-differentiated C57BL/6J WT Th17 cells treated with PI3K inhibitor (LY294002) for last 24 hours of differentiation. Data are mean±SEM (n=3). (F) Quantification of qPCR determination of IL-22 mRNA expression of *in vitro*-differentiated *Gclc^fl/fl^* and *Cd4Cre Gclc^fl/fl^* Th17 cells. Data are mean±SD (n=3-4). (G) IL-22 protein concentrations as measured by ELISA in culture supernatants of *in vitro*-differentiated *Gclc^fl/fl^* and *Cd4Cre Gclc^fl/fl^* Th17 cells. Data are mean±SEM (n=3); 2 trials. (H) Quantification of FCA of p-AKT in *in vitro-*differentiated *Gclc^fl/fl^* and *Cd4Cre-Gclc^fl/fl^* Th17 cells. Data are mean±SEM (n=3); 2 trials. (I) Left: FCA to detect p-mTOR in *in vitro-*differentiated *Gclc^fl/fl^* and *Cd4Cre-Gclc^fl/fl^* Th17 cells. Right: Quantification of left panel results. Data are mean±SEM (n=3); 3 trials. (J) Quantification of FCA of p-4E-BP1 in *in vitro-*differentiated *Gclc^fl/fl^* and *Cd4Cre-Gclc^fl/fl^* Th17 cells. Data are mean±SEM (n=3); 2 trials.

Mitochondrial gene expression is regulated by the mitochondrial transcription factor A (TFAM), which is crucial for OXPHOS (Desdín-Micó et al., 2020, Fu et al., 2019, Baixauli et al., 2015). In line with the reduced OXPHOS and the decreased expression of the mitochondrial encoded ETC genes, we also observed that expression of TFAM is downregulated in *Gclc*-deficient pathogenic Th17 cells, which can be restored by ROS-sacavenging by NAC (Figure 7B). This suggests the existence of a yet unappreciated feedback loop where mitochondrial respiration largely controls the generation of metabolic ROS (Dan Dunn et al., 2015), which when not scavenged by GSH negatively affects TFAM, mitochondrial ETC genes and decreases OXPHOS. In line with this ROS-induced negative mitochondrial cascade, ATP production was decreased in *Gclc*-deficient *in vitro*-differentiated pathogenic Th17 cells (Figure 7C). Again, ROS-scavenging by NAC could restore cellular ATP levels (Figure 7C). ATP is transported out of the mitochondria and fosters many cellular pathways. Especially, cellular signaling is dependent on ATP as it serves as a general kinase substrate. Consequently, low ATP concentrations can influence kinase activities depending on their affinity for ATP (Km-ATP).

We wondered whether the reduced ATP concentrations found in mutant Th17 cells and their inability to express IL-22 are linked. The Km-ATP of PI3Kδ, which is an upstream kinase in the mTOR activation pathway, is relatively high (118 mM), indicating that full activation of PI3K requires substantial amounts of ATP (Somoza et al., 2015). In line with the reduced ATP, we found that activation of PI3K was decreased in *Gclc-*deficient Th17 cells in a manner reversed by ROS-scavenging (Figure 7D). Moreover, specific inhibition of PI3K lead to downregulation of IL-22 production in the *in vitro*-differentiated WT pathogenic Th17 cells suggesting that these pathways might be linked (Figure 7E). Interestingly, we did not observe a downregulation of IL-22 encoding mRNA in mutant Th17 cells (Figure 7F). However, IL-22 protein expression was largely impacted in a ROS-dependent manner in *Gclc*-deficient pathogenic Th17 cells, ruling out a transcriptional regulation of this pathway (Figure 7G). On the contrary, IL-17 production was affected on both mRNA and protein levels (Figure S7G, H). This data suggests a ROS- and PI3K-dependent translational control of IL-22 production in Th17 cells. PI3K can stimulate protein translation by the PI3K/AKT/mTOR signaling pathway. In line, both phosphorylation of AKT and mTOR were significantly down-regulated in *Gclc-*deficient Th17 cells, which were restored by ROS-scavenging (Figure 7H-I). mTOR is not only crucial for host resistance to *C. rodentium* (Lin et al., 2016), but it also regulates translation. One mTOR target is the repressor of translation 4E-BP1. mTOR-dependent phosphorylation of 4E-BP1 induces the dissociation from translation initiation factor eIF4E, which then initiates protein translation (Sonenberg and Hinnebusch, 2009). In line with decreased mTOR and IL-22, 4E-BP1 phosphorylation was decreased in *Gclc-*deficient Th17 cells, while 4E-BP1 phosphorylation remained at control levels upon ROS-scavenging (Figure 7J).

Collectively, these results indicate that the reduced IL-22 production by *Gclc-*deficient Th17 cells is linked to reduced protein translation and mitochondrial function. Thus, physiological control of ROS is necessary for the generation of the IL-22-producing Th17 cells needed to confer protective immunity during GI infections. Taken together, our data reveal a T cell-intrinsic GSH-IL-22 signaling axis that is fundamental to the integrity of the gut intestinal barrier and depends on the control of mitochondrial ROS.

## Discussion

The GI tract is constantly exposed to microbial antigens of commensal and pathogenic organisms. Th cells in the gut maintain immune tolerance to the microbiome and mount immune responses against pathogens (van Wijk and Cheroutre, 2010). The GI tract is also a major site of ROS generated by food processing and immune cell interactions (Aviello and Knaus, 2017). Although low ROS support functions of T cells, elevated oxidative stress interferes with their effector actions (Devadas et al., 2002, Gülow et al., 2005, Jackson et al., 2004, Sena et al., 2013, Yi et al., 2006, Mak et al., 2017, Lang et al., 2013). Accordingly, T cells contain antioxidants such as GSH that scavenge intracellular ROS and limit their accumulation (Mak et al., 2017, Lian et al., 2018, Liang et al., 2006). In conventional T cells, GSH is essential for the metabolic reprogramming triggered by activation (Mak et al., 2017). In contrast, GSH-deficient Tregs show increased metabolic activity and are hyperproliferative but less suppressive (Kurniawan et al., 2020). Thus, GSH has T cell subset-specific functions that all contribute to immune homeostasis. The present study has revealed the role of GSH in Th17 cells, which are central effectors in mucosal immune responses. Loss of GSH-mediated ROS-buffering in Th17 cells compromises the IL-22 production needed to defend against bacterial GI infections.

Although several Th subsets contribute to immunity against *C. rodentium* including Th1, Th22 and Th17 cells, this pathogen elicits stronger Th17 responses (Higgins et al., 1999, Basu et al., 2012, Backert et al., 2014, Mangan et al., 2006). In line with this finding, we have shown that the dominant Th cells responding to *C. rodentium* infection are CD4^+^IL-17^+^IL-22^+^ Th17 cells. WT LP CD4^+^ T cells increased their GSH production in response to *C. rodentium* infection, suggesting a key role for GSH in these T cells. Consequently, *Gclc* deficiency had an important effect on IL-17^+^IL-22^+^ Th17 cells. Our *in vitro* and *in vivo* studies have shown that ablation of *Gclc* in Th17 cells is associated with the dysfunctionality of this important Th cell subset resulting in a fatal infection outcome.

Our work has demonstrated that mitochondria in *Gclc*-deficient LP Th17 cells, as well as mitochondria in *in vitro*-differentiated mutant Th17 cells, show compromised mitochondrial capacity, which has been shown to impair Th17 cell function (Kaufmann et al., 2019). Moreover, we observed dysregulation of expression of mitochondria-encoded ETC encoded genes and their transcription regulator TFAM. TFAM has been linked to the stabilization of mitochondrial DNA (mtDNA), its replication and OXPHOS (Ekstrand et al., 2004, Baixauli et al., 2015). In T cells, TFAM deletion has been shown to skew these cells towards a pro-inflammatory response (Desdín-Micó et al., 2020), specifically affecting immune surveillance by Tregs (Fu et al., 2019). Our data is line with these findings as mitochondrial dysfunction seems to affect mostly protective T cell responses, such as IL-22 production. Moreover, it has been suggested that TFAM in RORγT^+^ lymphocytes plays a key role in small intestine homeostasis (Fu et al., 2021). However, some questions still remain – is mROS dysregulation leading to mutation of mitochondria-encoded genes, leading to altered ETC activity as suggested for metastatic tumor cells (Ishikawa et al., 2008, Woo et al., 2012), or is the ROS-induced halt of mitochondrial transcription the sole mechanism leading to dysfunctional mitochondria? Our data, especially our immediate ROS-scavenging experiments suggest the latter could be the case.

Due to abnormal mitochondrial function, decreases in OCR and total ATP were observed in cultures of *Gclc*-deficient Th17 cells. Low ATP concentrations have recently been linked to decreased PI3K activation (Xu et al., 2021), and the PI3K/AKT/mTOR signaling pathway couples metabolic and transcriptional responses (Chi, 2012). We found that PI3K, AKT and mTOR activation were all decreased in mutant Th17 cells *in vitro,* in line with a previous report that oxidative stress inhibits PI3K/AKT/mTOR signaling in melanoma cells (Hambright et al., 2015). We therefore propose that a lack of ROS buffering in *Gclc*-deficient Th17 cells interferes with TFAM expression and transcription of mitochondrial encoded genes of the ETC. This leads to reduced OXPHOS and reduced mitochondrial ATP production resulting in lower cellular ATP concentrations associated with decreased PI3K/AKT/mTOR activation in Th17 cells. Activated mTORC1 enables release of the translation initiation factor eIF4E by phosphorylating the repressor of translation 4E-BP1 (Sonenberg and Hinnebusch, 2009). The the low 4E-BP1 phosphorylation in *Gclc*-deficient Th17 cells is indeed associated with reduced production of IL-22. Consistently, interference with mitochondrial translation has been previously connected with T cell cytokine production (Almeida et al., 2021, Colaço et al., 2021). *In vitro* as well as *in vivo,* we have shown that ROS-accumulation in Th17 cells is decisive for the mitochondrial dysfunction and impaired cytokine production. Accordingly, the lethality of *C. rodentium-*infected *Cd4^Cre^-Gclc^fl/fl^* mice could be prevented by *in vivo* ROS-scavenging.

Although CD4-expressing LTi cells also co-produce IL-22 and IL-17, we found no impact of *Gclc* deletion on cytokine production by caecal or colonic LTi cells during *C. rodentium* infection. In addition, the course of *C. rodentium* infection was nearly identical in *Rag1^-/-^ Cd4Cre Gclc^fl/fl^* and control *Rag1^-/-^Gclc^fl/fl^* mice, indicating that loss of *Gclc* in LTi cells has no effect on the severe outcomes of infection in *Cd4Cre Gclc^fl/fl^* mice. Intracellular ROS levels in LTi cells were unaffected by *Gclc* deletion, suggesting that, unlike in T cells, LTi cell function does not depend on *Gclc*-mediated ROS buffering. Indeed, mitochondrial metabolism and the impact of mitochondrial ROS in ILC3s are known to diverge from that in Th17 cells. Whereas Th17 cells maintain low levels of mitochondrial ROS to avoid detrimental effects, high mitochondrial ROS promote ILC3 effector function, and antioxidants impair ILC3 cytokine production (Di Luccia et al., 2019). Consistent with this report, we found that intracellular ROS were much higher in T cells than in LTi cells after *Gclc* ablation (Figure 3D, S3F). Lastly, unlike T cells, ILCs and other innate immune cells could rely more on alternative antioxidants to compensate for loss of GSH-mediated ROS-buffering capacity (Yang et al., 2020).

In line with the finding that only mice lacking IL-22 are unable to clear *C. rodentium* infections (Basu et al., 2012, Zheng et al., 2008, Ota et al., 2011, Ishigame et al., 2009, Guo et al., 2014), we showed that IL-22-Fc treatment *in vivo* protected infected *Cd4Cre Gclc^fl/fl^* mice from lethality and that genetic restoration of T cell-mediated IL-22 production, but not IL-17, in *Cd4Cre Gclc^fl/fl^* mice allowed them to clear the pathogen. Although IL-22 controls AMP expression (Liang et al., 2006, Zheng et al., 2008, Raffatellu et al., 2009, Dixon et al., 2016), we saw no differences in these peptides between mutant and control mice. It may be that ILC-derived IL-22 induces adequate AMP expression in *Gclc*-deficient mice. However, it is T cell-derived IL-22 that is essential to regulate tight junction integrity and mucus production, which protects the intestine from *C. rodentium*-induced damage during the late stages of *C. rodentium* infection (Zindl et al., 2021). Accordingly, infected *Cd4Cre Gclc^fl/fl^* mice exhibited severely disorganized tight junctions, reduced IEC MUC2 content and a thin mucus layer. As a result, gut permeability was increased in the mutants, allowing bacterial spread to the peripheral organs and blood. Injection of IL-22-Fc preserved normal tight junctions, IEC MUC2 content and a robust mucus layer in infected mutants. Pertinently, loss of intestinal barrier function, reduced mucin production, increased oxidative stress and decreased antioxidant capacity have all been linked to chronic inflammation in IBD patients (Shorter et al., 1972, Corridoni et al., 2014, Sido et al., 1998, Buffinton and Doe, 1995, Lih-Brody et al., 1996, Kruidenier et al., 2003). In line, we saw a positive correlation between *GCLC* expression and epithelial integrity markers in rectal tissue sections of UC patients. The expression of Th17 cell markers correlated positively with *GCLC* expression, hinting that *GCLC* regulates Th17 cell functions and thus intestinal epithelial integrity in humans with GI pathology. Our results provide a rationale for exploring antioxidant treatment as a strategy to regulate Th17-derived IL-22 production in these disorders.

In conclusion, our findings imply the existence of a previously unappreciated axis between GSH, mitochondrial function and IL-22 signaling within Th17 cells that operates during GI infections. If ROS accumulate in Th17 cells, TFAM, mitochondrial activity and PI3K/AKT/mTOR are dysregulated, impairing IL-22 protein production. IL-22 is critical for intestinal barrier integrity and mouse survival during bacterial GI infection. Further study of this newly described GSH-dependent regulatory circuit in LP Th17 cells may yield new insights into how increased oxidative stress affects the function of this T cell subset and leads to intestinal pathology. Our results may also point to novel therapeutic strategies for modulating Th17 cell function in the context of human GI disorders.

## Acknowledgements

Aymeric Fouquier d’Herouël (LCSB, Luxembourg) and Olga Kondratyeva (LCSB, Luxembourg) for support in FACS sorting; Dominique Revets and the National Cytometry Platform (LIH, Luxembourg) for support in immunofluorescence microscopy and flow cytometry; and Christoph Wilhelm (Univ. Bonn) for technical advice. We are also grateful to Samantha Storn, Anaïs Oudin, all Animal Facility staff, and LIH’s Animal Welfare Structure for animal services at LIH, Luxembourg; and Djalil Coowar, Jennifer Behm; Marthe Schmit and all Animal Facility staff for animal services at Univ. Luxembourg.

D.B. is supported by FNR-ATTRACT (A14/BM/7632103) and FNR-CORE grants (C21/BM/15796788), (C18/BM/12691266). D.B., T.K. and C.D. are supported by FNRS-Televie grants No. 7.4587.20 and/or No. 7.497.19. D.B., L.B., L.G. and L.S-B. by FNR-PRIDE (PRIDE/11012546/NEXTIMMUNE); D.B. and A.E. by (PRIDE17/11823097/MicrOH); D.B. and D.G.F. by FNR-RIKEN (TregBar/11228353), and M.M. by FNR PEARL P16/BM/11192868. V.V. holds grant NIH/NIAAA (5R24AA022057). E.L. is supported by FNR-CORE grants (C16/BM/11282028 and C20/BM/14591557) and FNR-JUMP PoC/18/12554295 (E.L.) and FNR-MFP20/15251414. E.K. is supported by FNR-PoC/18/12554295. I.S.H. is supported by the American Association for Cancer Research and Breast Cancer Research Foundation (20-20-26-HARR). A.W. is supported by the German Research Foundation (DFG) under the project numbers 490846870 – TRR355/1 TPA08 and TPB05. P.A.L. is supported by the DFG (RTG1949, LA2558/8-1), the Volkswagen Foundation and the Jürgen Manchot Foundation (Molecules Of Infection).

## Author Contributions

L.B. performed most experiments assisted by V.H., L.G., H.K, D.G.F., L.S-B., M.G., A.E, C.B., C.V., and S.F. Lamina propria isolations were carried out by L.B. and L.G., H.K., L.B., L.S-B. and D.G.F. performed Seahorse flux assays. RNA-sequencing data analysis was performed by E.K. and J.L. and guided by E.L. J-J.G. performed H&E and Alcian Blue histology stainings. IL-22^ind^ mice were generated by S.S. and B.B. T.K., C.D., Y.C., V.V, B.B., I.S.H., P.A.L, A.W., and M.M. provided reagents and expert comments. D.B. supervised the study. L.B., and D.B. conceptualized the work, together with V.H. designed all experiments, analyzed the data, and wrote the manuscript. All authors reviewed and edited the final manuscript.

## Declaration of Interests

The authors declare no competing interests.

## Supplementary Figure Legends

**Supplementary Figure 1:**
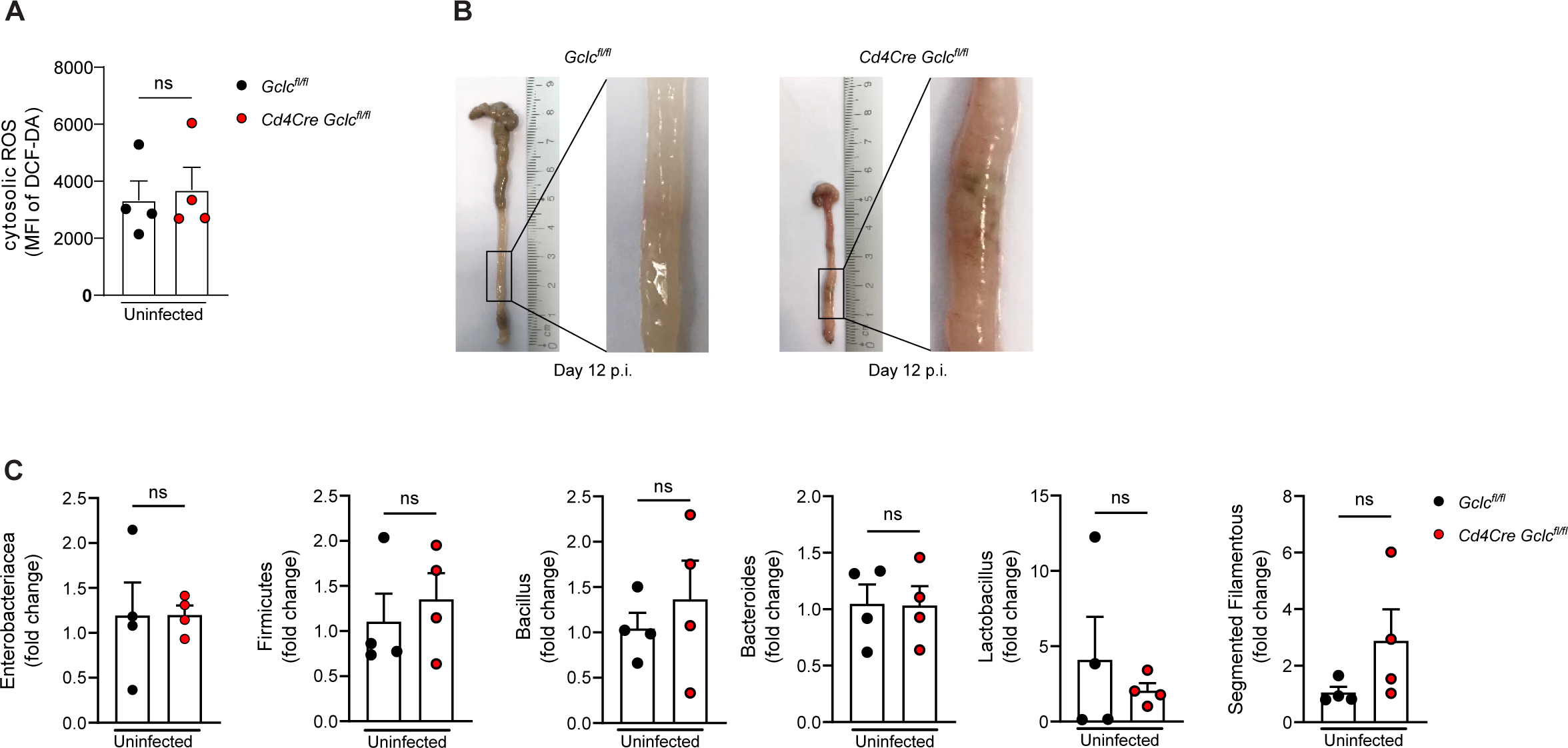
T cell-specific ablation of *Gclc* does not affect the gut microbiome. (A) Quantification of flow cytometric analysis of DCF-DA staining to detect intracellular ROS in CD4^+^ cells of the colonic LP, which was isolated from uninfected *Gclc^fl/fl^* and *Cd4Cre-Gclc^fl/fl^* mice. MFI, mean fluorescence intensity. Data are mean±SEM (n=3); 2 trials. (B) Macroscopic views and lengths of colons from *C. rodentium*-infected *Gclc^fl/fl^* and *Cd4Cre-Gclc^fl/fl^* mice at day 12 p.i. Images are representative of 5 mice per genotype; 2 trials. (C) Quantification of qPCR determinations of DNA of the indicated bacterial genera in feces of uninfected *Gclc^fl/fl^* and *Cd4Cre-Gclc^fl/fl^* mice. ΔΔCt values were normalized to total Eubacteria. Data are mean±SEM (n=4); 2 trials.

**Supplementary Figure 2:**
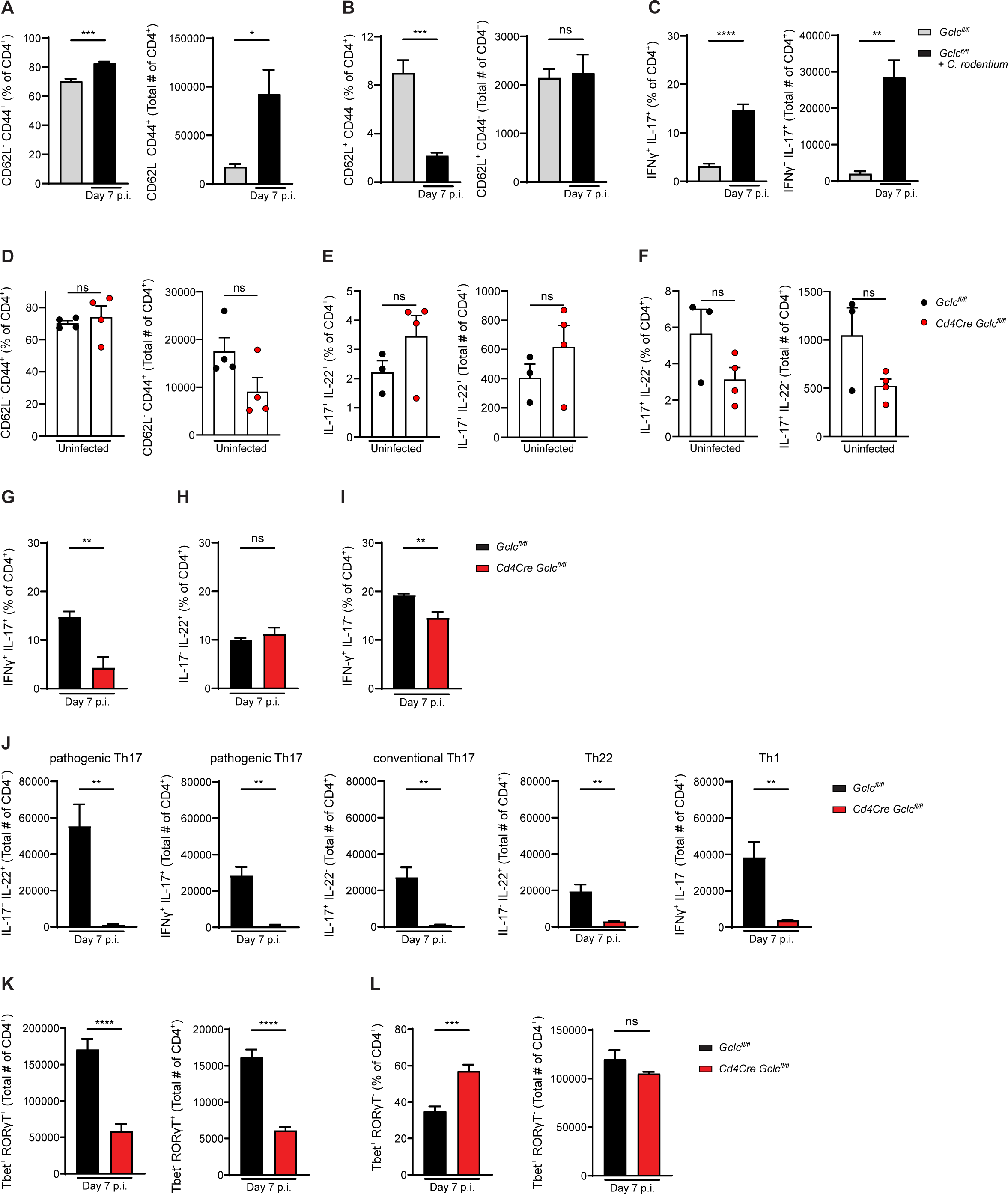
T cell-specific ablation of *Gclc* does not alter steady-state levels of IL-22 and IL-17 in colonic LP T cells. (A) Frequencies (left) and total numbers (right) of CD62L^-^CD44^+^ cells as determined by flow cytometric analysis of CD4^+^ cells of the colonic LP, which was isolated from uninfected and *C. rodentium*-infected *Gclc^fl/fl^* mice at day 7 p.i. Data are mean±SEM (n=4); 2 trials. (B) Frequencies (left) and total numbers (right) of CD62L^+^CD44^-^ cells as determined by flow cytometric analysis of CD4^+^ cells of the colonic LP, which was isolated as in (A). Data are mean±SEM (n=4); 2 trials. (C) Frequencies (left) and total numbers (right) of IFN-γ^+^IL-17^+^ cells as determined by flow cytometric analysis of CD4^+^ cells of the colonic LP, which was isolated as in (A). LP cells were stimulated with PMA/calcium ionophore/Brefeldin A for 5 hr before staining. Data are mean±SEM (n=4); 3 trials. (D) Frequencies (left) and total numbers (right) of CD62L^-^CD44^+^ cells as determined by flow cytometric analysis of colonic LP CD4^+^ cells isolated from uninfected *Gclc^fl/fl^* and *Cd4Cre-Gclc^fl/fl^* mice. Data are mean±SEM (n=4); 2 trials. (E) Frequencies (left) and total numbers (right) of IL-17^+^IL-22^+^ cells as determined by intracellular flow cytometric analysis of colonic LP CD4^+^ cells isolated as in (D). LP cells were stimulated with PMA/calcium ionophore/Brefeldin A for 5 hr before staining. Data are mean±SEM (n=4); 3 trials. (F) Frequencies (left) and total numbers (right) of IL-17^+^IL-22^-^ cells as determined by intracellular flow cytometric analysis of colonic LP CD4^+^ cells isolated as in (D). LP cells were stimulated as in (E). Data are mean±SEM (n=4); 3 trials. (G) Frequencies of IFN-γ^+^IL-17^+^ cells as determined by flow cytometric analysis of CD4^+^ cells of the colonic LP, which was isolated from *C. rodentium*-infected *Gclc^fl/fl^* and *Cd4Cre-Gclc^fl/fl^* mice at day 7 p.i. LP cells were stimulated with PMA/calcium ionophore/Brefeldin A for 5 hr before staining. Data are mean±SEM (n=4); 3 trials. (H) Frequencies of IL-17^-^IL-22^+^ cells as determined by flow cytometric analysis of CD4^+^ cells of the colonic LP, which was isolated as in (G). LP cells were stimulated as in (G). Data are mean±SEM (n=4); 3 trials. (I) Frequencies of IFN-γ^+^IL-17^-^ cells as determined by flow cytometric analysis of CD4^+^ cells of the colonic LP, which was isolated and treated as in (G). LP cells were stimulated as in (G). Data are mean±SEM (n=4); 3 trials. (J) Total cell numbers as determined by intracellular flow cytometric analysis of the indicated CD4^+^ T cell subsets among colonic LP cells, which were isolated from *C. rodentium*-infected *Gclc^fl/fl^* and *Cd4Cre-Gclc^fl/fl^* mice at day 7 p.i. LP cells were stimulated as in (G). Data are mean±SEM (n=4); 3 trials. (K) Total cell numbers as determined by intracellular flow cytometric analysis of CD4^+^Tbet^+^RORγT^+^ (left) and CD4^+^Tbet^-^RORγT^+^ (right) cells of the colonic LP, which was isolated from *C. rodentium*-infected *Gclc^fl/fl^* and *Cd4Cre-Gclc^fl/fl^* mice at day 7 p.i. Data are mean±SEM (n= 6); 2 trials. (L) Frequencies (left) and total numbers (right) of Tbet^+^RORγT^-^ cells as determined by intracellular flow cytometric analysis of CD4^+^ cells in the colonic LP, which was isolated as in (K). Data are mean±SEM (n= 6); 2 trials.

**Supplementary Figure 3:**
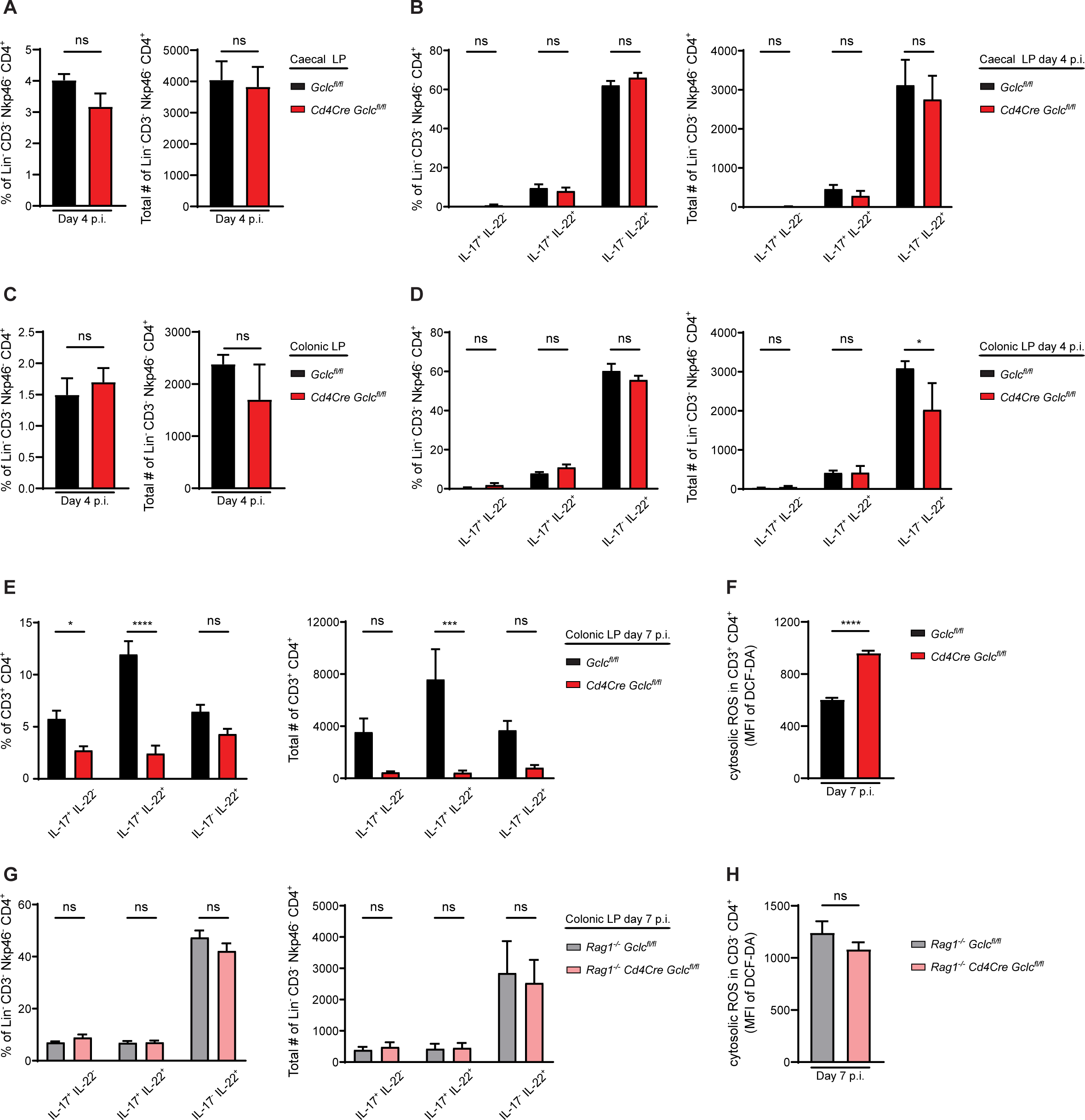
*Gclc* regulates IL-17 and IL-22 expression in T cells but not in LTi cells. (A) Quantification of flow cytometric analysis of frequencies (left) and total numbers (right) of LTi cells in caecal LP isolated from *C. rodentium*-infected *Gclc^fl/fl^* and *Cd4Cre-Gclc^fl/fl^* mice at day 4 p.i. Data are mean±SEM (n=4-5). (B) Quantification of flow cytometric analysis of frequencies (left) and total numbers (right) of the indicated LTi subpopulations from the data in (A) at day 4 p.i. LP cells were stimulated with PMA/calcium ionophore/Brefeldin A for 5 hr before staining. Data are mean±SEM (n=4-5). (C) Frequencies (left) and total cell numbers (right) as determined by flow cytometric analysis of total LTi cells among colonic LP cells isolated from *C. rodentium*-infected *Gclc^fl/fl^* and *Cd4Cre-Gclc^fl/fl^* mice at day 4 p.i. Data are mean±SEM (n=4-5). (D) Frequencies (left) and total cell numbers (right) as determined by intracellular flow cytometric analysis of the indicated LTi subpopulations among colonic LP cells isolated as in (C). LP cells were stimulated as in (B). Data are mean±SEM (n=4-5). (E) Frequencies (left) and total cell numbers (right) of the indicated populations as determined by flow cytometric analysis of intracellular IL-17 and IL-22 staining in CD3^+^CD4^+^ T cells among colonic LP cells of *C. rodentium*-infected *Gclc^fl/fl^* and *Cd4Cre-Gclc^fl/fl^* mice at day 7 p.i. LP cells were stimulated as in (B). Data are mean±SEM (n=5); 3 trials. (F) Quantification of flow cytometric analysis of intracellular ROS in CD3^+^CD4^+^ T cells of the colonic LP, which was isolated from *C. rodentium*-infected *Gclc^fl/fl^* and *Cd4Cre-Gclc^fl/fl^* mice at day 7 p.i. and subjected to DCF-DA staining. Data are mean±SEM (n=4-5). (G) Frequencies (left) and total cell numbers (right) as determined by intracellular flow cytometric analysis of the indicated subpopulations of LTi cells among colonic LP cells isolated from *C. rodentium*-infected *Rag1^-/-^Gclc^fl/fl^* and *Rag1^-/-^Cd4Cre-Gclc^fl/fl^* mice at day 7 p.i. LP cells were stimulated as in (B). Data are mean±SEM (n=5). (H) Quantification of flow cytometric analysis of intracellular ROS in CD3^-^CD4^+^ LTi cells of the colonic LP, which was isolated as in (G) and subjected to DCF-DA staining. Data are mean±SEM (n=5).

**Supplementary Figure 4:**
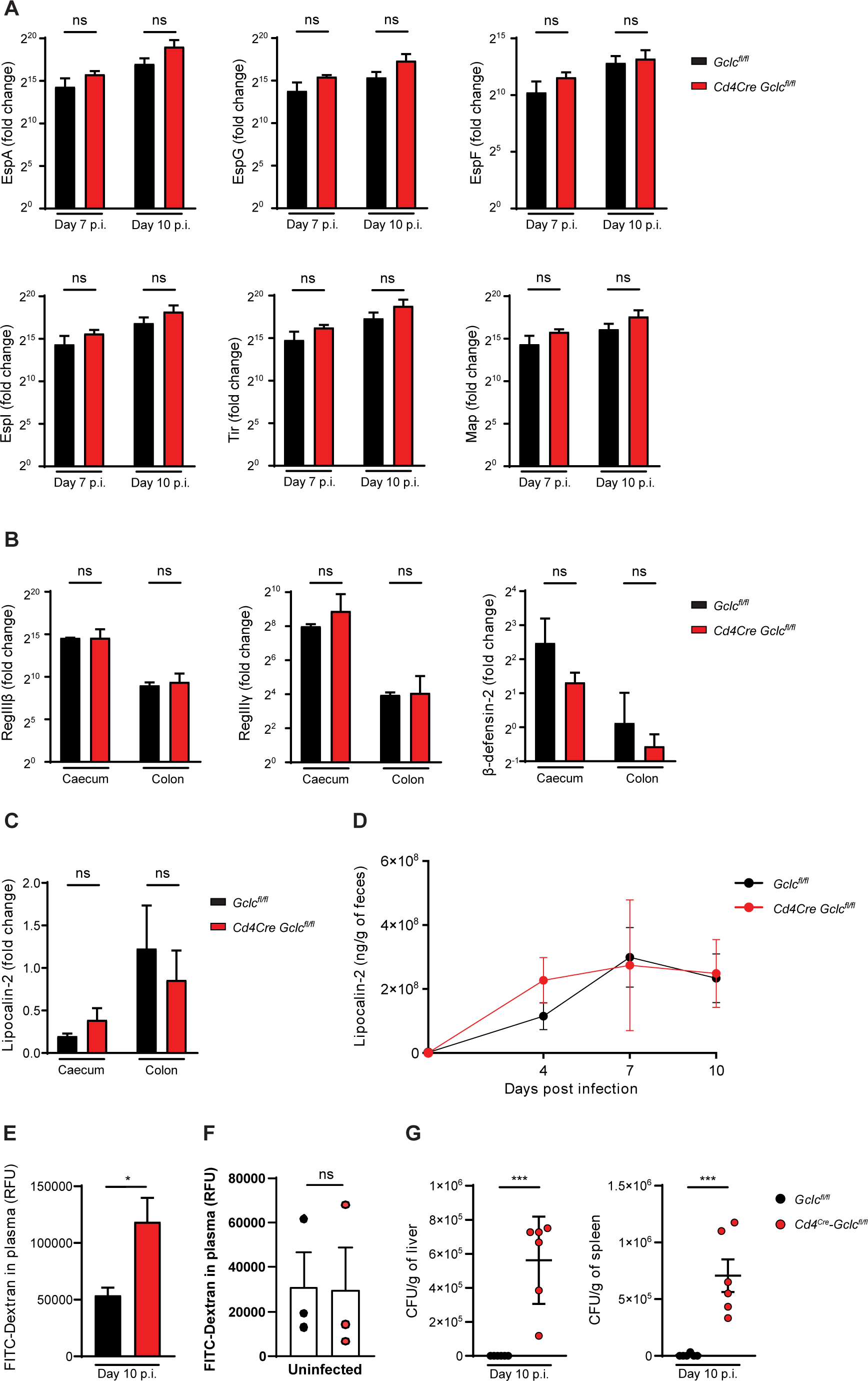
Ablation of *Gclc* in T cells does not affect either bacterial virulence factors of *C. rodentium* or intestinal anti-microbial peptides (AMPs). (A) RT-qPCR determination of mRNA levels of the indicated bacterial virulence factors in colonic tissues of *C. rodentium*-infected *Gclc^fl/fl^* and *Cd4Cre-Gclc^fl/fl^* mice at day 7 and 10 p.i. ΔΔCt values were normalized to Rps17 expression. Data are mean±SEM (n=3-4); 2 trials. (B) RT-qPCR determination of mRNA levels of the indicated AMPs in caecal or colonic tissues of *C. rodentium*-infected *Gclc^fl/fl^* and *Cd4Cre-Gclc^fl/fl^* mice at day 7 p.i. ΔΔCt values were normalized to Hprt expression. Data are mean±SEM (n=3-4); 3 trials. (C) RT-qPCR determination of mRNA levels of lipocalin-2 in caecal or colonic tissues of *C. rodentium*-infected *Gclc^fl/fl^* and *Cd4Cre-Gclc^fl/fl^* mice at day 10 p.i. ΔΔCt values were normalized to Hprt expression. Data are mean±SEM (n=4); 2 trials. (D) ELISA determinations of lipocalin-2 protein in supernatants of fecal homogenates of *C. rodentium*-infected *Gclc^fl/fl^* and *Cd4Cre-Gclc^fl/fl^* mice collected at the indicated time points after infection. Data are mean±SEM (n=4); 3 trials. (E) Quantification of plasma FITC-Dextran levels in *C. rodentium*-infected *Gclc^fl/fl^* and *Cd4Cre-Gclc^fl/fl^* mice that were gavaged with FITC-Dextran at day 10 p.i. FITC-Dextran was measured 4 hr later. Data are mean±SEM (n=4-5); 3 trials. (F) Quantification of plasma FITC-Dextran levels in uninfected *Gclc^fl/fl^* and *Cd4Cre-Gclc^fl/fl^* mice that were gavaged with FITC-Dextran. FITC-Dextran was measured 4 hr later. Data are mean±SEM (n=3). (G) CFU of *C. rodentium* in liver (left) and spleen (right) of *C. rodentium*-infected *Gclc^fl/fl^* and *Cd4Cre-Gclc^fl/fl^* mice at day 10 p.i. Data are mean±SEM (n=6); 3 trials.

**Supplementary Figure 5:**
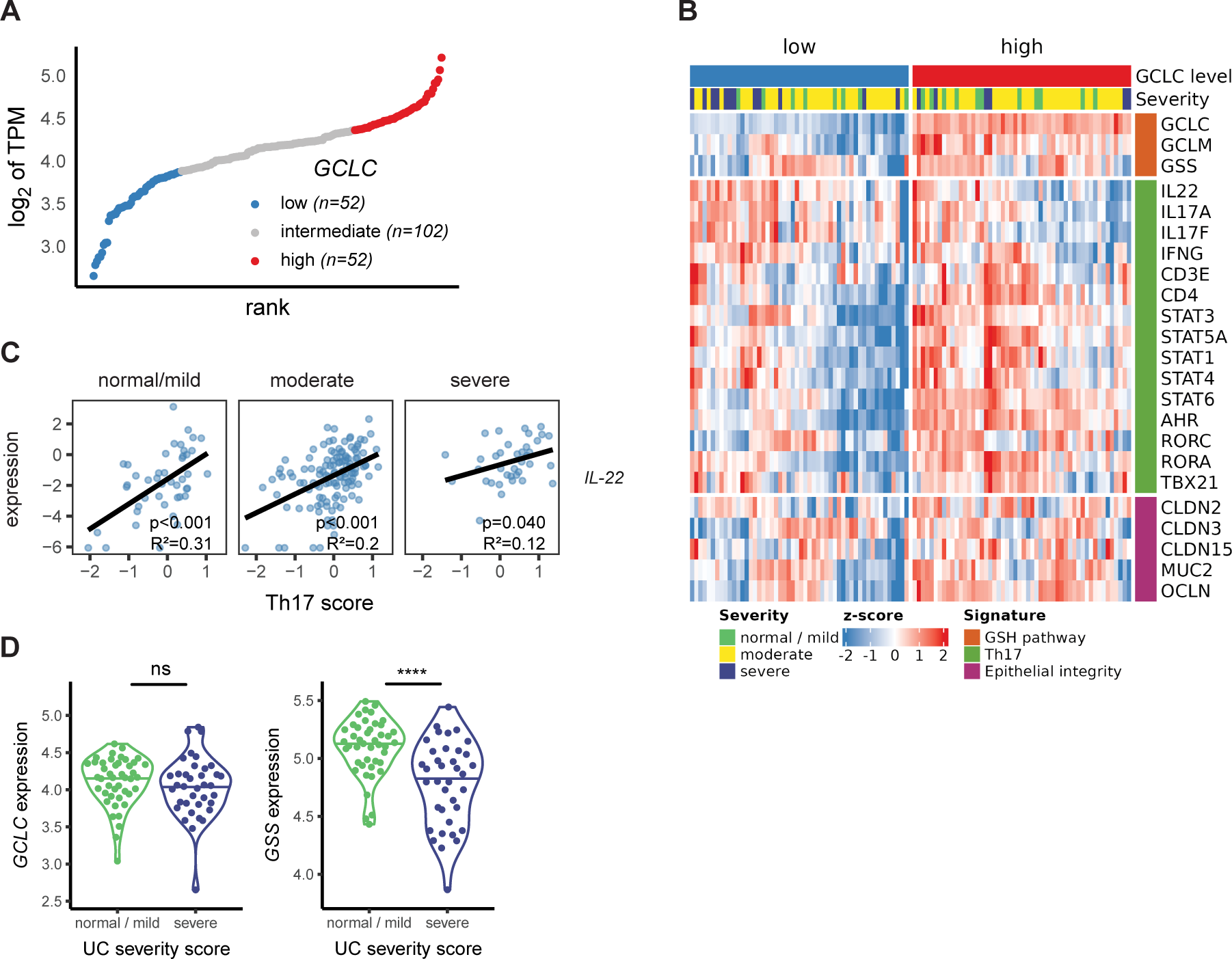
*GSS* expression is decreased in rectal tissues of IBD patients with severe pathology. (A) Bioinformatics analysis of RNA sequencing data of rectal biopsies from ulcerative colitis (UC) patients (Haberman et al., 2019). The lower and upper quartile values of *GCLC* expression were used to classify the patients into low, intermediate and high expressors. (B) Heat-map representation of selected signature genes of interest in the low and high *GCLC*-expressing UC patients in (A). The provided MAYO scores were used to define the disease severity as normal/mild, moderate and severe disease (Teixeira et al., 2015). (C) Bioinformatics analysis of RNA sequencing data of rectal biopsies from the UC patients in (A,B) examining the correlation between Th17 score and *IL-22* expression. Significance and r^2^ values are indicated in the lower right corner of each frame. The Th17 score was defined as the mean of scaled log2 gene expression values composing the signature (*IL-17A, IL-17F, IFN-γ, CD3E, CD4, STAT3, STAT5a, STAT1, STAT4, STAT6, AHR, RORC, RORA and TBX21*). (D) Violin plots of *GCLC* (left) and *GSS* (right) expression according to UC disease severity (showing normal/mild versus severe scores) from the data in (A). Significance was assessed using the Wilcoxon signed rank test.

**Supplementary Figure 6:**
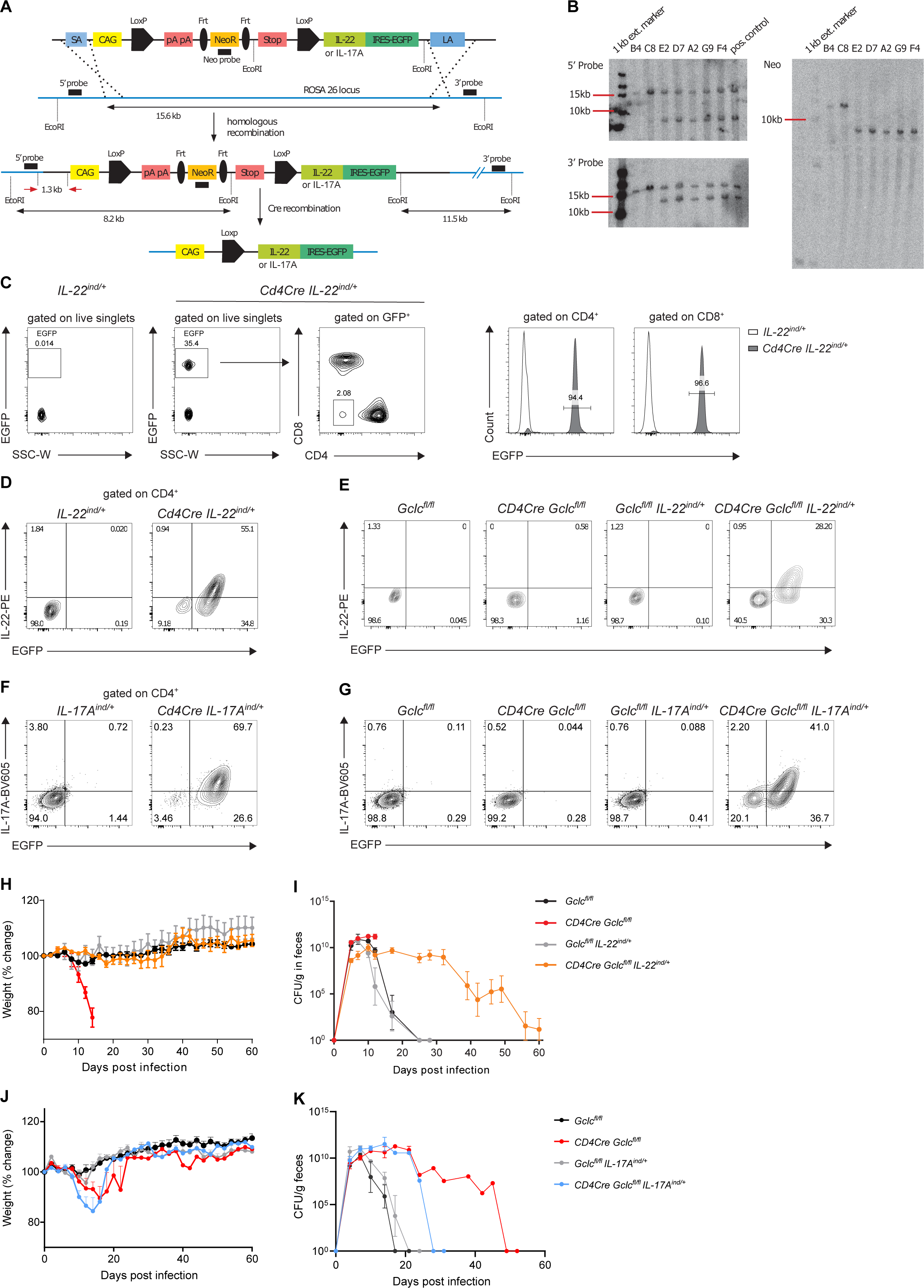
T cell-derived IL-22 is sufficient for mice with GSH-deficient T cells to survive and clear *C. rodentium* infection. (A) Schematic representation of the vector bearing the mouse *IL-22* (or *IL-17)* and *EGFP* genes into the *Rosa26* locus by homologous recombination. Upon Cre recombinase-mediated excision of the stop cassette, IL-22 (or IL-17) and EGFP are expressed under the CAG promotor. Southern blotting probes and fragments (after digestion with EcoRI) are depicted in black, with screening PCR primers depicted in red. SA/LA, short/long homology arm; CAG, CMV early enhancer/chicken β actin promoter; pA, poly A; stop, Westphal stop sequence; NeoR, neomycin resistance; EGFP, enhanced green fluorescent protein; ES cells, embryonic stem cells. (B) Southern blot analyses of 7 positive ES cell clones identified by PCR using the external 5’ and 3’ probes and the Neo probe indicated in (A). WT bands are 15.6 kb for both probes, targeted bands are 8.2 or 11.5 kb for 5’ and 3’ probe, respectively, and 8.2 kb for the neo probe. (C) Flow cytometric analysis of EGFP expression by total live cells and by live CD4^+^ and CD8^+^ cells isolated from spleens of naïve *IL-22^ind/+^* and *Cd4Cre IL-22^ind/+^* mice. Plots are representative of 3 mice per genotype. (D) Flow cytometric analysis of EGFP and IL-22 expression by live CD4^+^ cells isolated from spleens of naïve *IL-22^ind/+^* and *Cd4Cre IL-22^ind/+^* mice. Cells were activated overnight by culture with soluble anti-CD3/anti-CD28 (5µg/mL) and restimulated with PMA/calcium ionophore/Brefeldin A for 5 hr before cytokine staining. Plots are representative of 3 mice per genotype. (E) Flow cytometric analysis of EGFP and IL-22 expression by live CD4^+^ cells isolated from spleens of naïve *Gclc^fl/fl^*, *Cd4Cre Gclc^fl/fl^*, *Gclc^fl/fl^IL-22^ind/+^* and *Cd4Cre Gclc^fl/fl^IL-22^ind/+^* mice. Cells were activated by culture overnight with plate-bound anti-CD3 (5µg/mL) and soluble anti-CD28 (1µg/mL) and restimulated for cytokine staining as in (D). Plots are representative of 3 mice per genotype. (F) Flow cytometric analysis of EGFP and IL-17A expression by live CD4^+^ cells isolated from spleens of naïve *IL-17A^ind/+^* and *Cd4Cre IL-17A^nd/+^* mice. Cells were activated overnight by culture with soluble anti-CD3/anti-CD28 (5µg/mL) and restimulated with PMA/calcium ionophore/Brefeldin A for 5 hr before cytokine staining. Plots are representative of 3 mice per genotype. (G) Flow cytometric analysis of EGFP and IL-17A expression by live CD4^+^ cells isolated from spleens of naïve *Gclc^fl/fl^*, *Cd4Cre Gclc^fl/fl^*, *Gclc^fl/fl^IL-17A^ind/+^* and *Cd4Cre Gclc^fl/fl^IL-17A^ind/+^* mice. Cells were activated by culture overnight with plate-bound anti-CD3 (5µg/mL) and soluble anti-CD28 (1µg/mL) and restimulated for cytokine staining as in (D). Plots are representative of 3 mice per genotype. (H) Change in whole body weight of *Gclc^fl/fl^* (n=5), *Cd4Cre Gclc^fl/fl^* (n=6)*, Gclc^fl/fl^*IL-22^ind/+^ (n=5) and *Cd4Cre Gclc^fl/fl^*IL-22^ind/+^ (n=8) mice infected with *C. rodentium* on day 0 and assayed on the indicated days p.i. Data are mean±SEM (n=5-8). (I) CFU of *C. rodentium* in feces of *Gclc^fl/fl^*, *Cd4Cre Gclc^fl/fl^*, *Gclc^fl/fl^IL-22^ind/+^* and *Cd4Cre Gclc^fl/fl^IL-22^ind/+^* mice. Mice were infected with *C. rodentium* on day 0 and feces were collected for analysis on the indicated days. Data are mean±SEM (n=5-8). (J) Change in whole body weight of *Gclc^fl/fl^* (n=6), *Cd4Cre Gclc^fl/fl^* (n=9)*, Gclc^fl/fl^IL-17A^ind^*^/+^ (n=6) and *Cd4Cre Gclc^fl/fl^IL-17A^ind^*^/+^ (n=7) mice infected with *C. rodentium* on day 0 and assayed on the indicated days p.i. Data are mean±SEM (n=6-9). (K) CFU of *C. rodentium* in feces of *Gclc^fl/fl^*, *Cd4Cre Gclc^fl/fl^*, *Gclc^fl/fl^IL-17A^ind/+^* and *Cd4Cre Gclc^fl/fl^IL-17A^ind/+^* mice. Mice were infected with *C. rodentium* on day 0 and feces were collected for analysis on the indicated days. Data are mean±SEM (n=6-9).

**Supplementary Figure 7:**
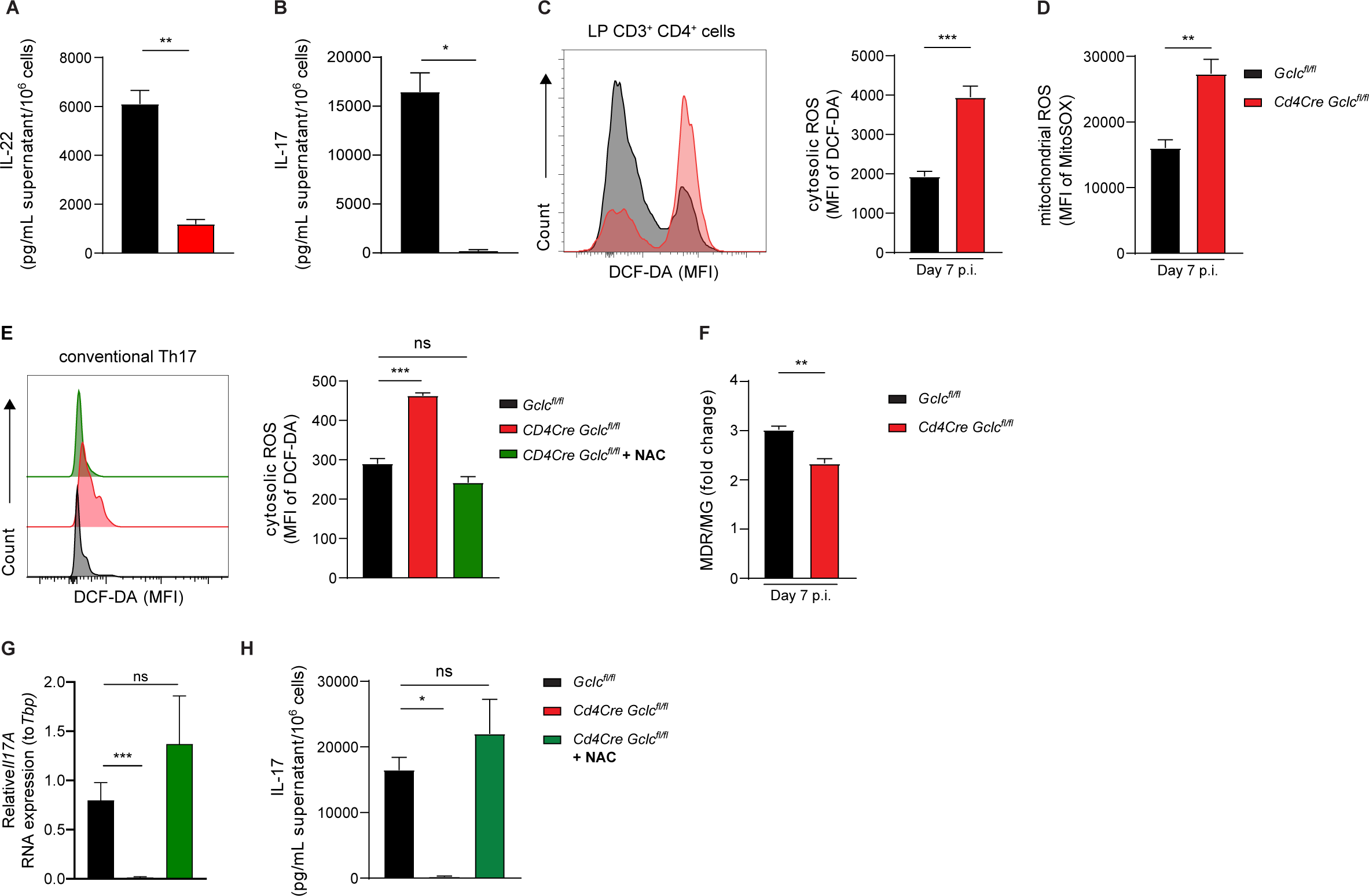
*Gclc* regulates mitochondrial function, ROS and IL-17. (A-B) Naïve T cells were sorted from spleen and lymph nodes of *Gclc^fl/fl^* and *Cd4Cre Gclc^fl/fl^* mice and induced to differentiate *in vitro* into pathogenic Th17 cells by culture with anti-CD3, anti-CD28, IL-6, IL-1β and IL-23. (A) IL-22 and (B) IL-17 were quantified by ELISA. Data are mean±SEM (n=3); 2 trials. (C) Left: Flow cytometric analysis of DCF-DA staining to detect intracellular ROS in CD3^+^CD4^+^ cells of the colonic LP, which was isolated from *C. rodentium*-infected *Gclc^fl/fl^* and *Cd4Cre Gclc^fl/fl^* mice at day 7 p.i. Right: Quantification of the data in the left panel. Data are mean±SEM (n=4). (D) Quantification of MitoSOX staining to detect mitochondrial ROS in CD3^+^CD4^+^ cells of the colonic LP, which was isolated from *C. rodentium*-infected *Gclc^fl/fl^* and *Cd4Cre Gclc^fl/fl^* mice at day 7 p.i. Data are mean±SEM (n=4). (E) Naïve CD4^+^ T cells were sorted from spleen and lymph nodes of *Gclc^fl/fl^* and *Cd4Cre Gclc^fl/fl^* mice and induced to differentiate *in vitro* into conventional Th17 cells by culture with anti-CD3, anti-CD28, IL-6, and TGF-β. Cells were treated with 10mM NAC as indicated. Left: Flow cytometric analysis of DCF-DA to detect ROS in *Gclc^fl/fl^* and *Cd4Cre Gclc^fl/fl^ in vitro-* differentiated conventional Th17 cells. Right: Quantification of the data in the left panel. Data are mean±SEM (n=3); 2 trials. (F) Determination of MDR/MG ratio to assess mitochondrial activity per mitochondrial mass in the cells in (A, B). Data are mean±SEM (n=4). (G) Quantification of qPCR determination of IL-17A mRNA expression of in vitro-differentiated *Gclc^fl/fl^* and *Cd4Cre Gclc^fl/fl^* pathogenic Th17 cells. Data are mean±SD (n=3-4). (H) IL-17A protein concentrations as measured by ELISA in culture supernatants of *in vitro*-differentiated *Gclc^fl/fl^* and *Cd4Cre Gclc^fl/fl^* pathogenic Th17 cells. Data are mean±SEM (n=3); 2 trials.

## Materials and Methods

### Mice

*Cd4Cre Gclc^fl/fl^* mice have been previously described (Mak et al., 2017). *Rag1^-/-^Cd4Cre Gclc^fl/fl^* mice were obtained by crossing *Cd4Cre Gclc^fl/fl^* with *Rag1^-/-^* mice. *Rag1^-/-^* and C57BL/6J mice were purchased from The Jackson Laboratory, *IL-17A^ind^* mice (Haak et al., 2009) were provided by Ari Waisman (Mainz, Germany). All mice were bred in the specific pathogen-free (SPF) facility of the Luxembourg Institute of Health (LIH).

*IL-22^ind^* mice [C57BL6/J-Gt(ROSA)26Sor<tm1(IL-22-ind)TgFl] were generated by Sabine Spath in the laboratory of Burkhard Becher at the Institute of Experimental Immunology (University of Zurich, Switzerland). The *Rosa26* targeting vector containing homology arms for the *Rosa26* locus was generated by Tobias Heinen in the laboratory of Ari Waisman (Mainz, Germany). The cDNA containing *IL-22* and *EGFP* was integrated by conventional cloning and is preceded by a loxP-flanked stop cassette which, when removed by Cre-mediated recombination, allows expression of IL-22 and EGFP from the upstream CAG promotor. The verified targeting vector was linearized and purified for injection using phenol/chloroform extraction.

Homologous recombination and targeting of the *Rosa26* locus were performed by the transgenic facility of the B.S.R.C. “Alexander Fleming” in Greece using conventional targeting by electroporation in Bruce4 C57BL/6 ES cells. ES cells were screened by PCR for integration of the construct: primer forward, external *Rosa26* (TAG GTA GGG GAT CGG GAC TCT), primer reverse, mutant *Rosa26* (GCG AAG AGT TTG TCC TCA ACC). Using the Taq PCR Core Kit (Qiagen), the following touchdown PCR was performed: 2 cycles each at 60°C, 59°C and 58°C, then 35 cycles at 57°C. A band at 1300 bp indicated homologous recombination. Correct integration was further confirmed by Southern blot (c.f. Figure S6B) in which genomic DNA was digested with EcoRI and separated on a 0.7% agarose gel. The following probes were used: *Rosa26* external 5’ probe ‘Orkin’ (restriction digest with EcoRI and PacI, 698 bp), modified from (Mao et al., 1999) and *Rosa26* external 3’ probe ‘Soriano’ (restriction digest with EcoRI, 700 bp), adapted from (Awatramani et al., 2001, Soriano, 1999). Probes (50 ng) were labeled with ^32^P using the Ladderman DNA labeling kit (TaKaRa) according to the manufacturer’s protocol, and Southern blotting was performed according to standard protocols.

Clone E2 was selected for downstream procedures. Chimeras were generated by injecting targeted ES cells into C57BL/6 albino mouse blastocysts. Routine genotyping of *IL-22^ind^* mice was performed using the following primers at 58°C: Primer *Rosa26* FW (AAA GTC GCT CTG AGT TGT TAT), Primer *Rosa26* RW (GGA GCG GGA GAA ATG GAT ATG), Primer SpliceAcB (CAT CAA GGA AAC CCT GGA CTA CTG). The WT band appeared at 600 bp and the recombinant band at 250 bp.

*Cd4Cre IL-22^ind^* mice were obtained by crossing *IL-22^ind^* mice with *Cd4Cre* mice purchased from The Jackson Laboratory*. Cd4Cre Gclc^fl/fl^IL-22^ind^* mice were generated at the Luxembourg Institute of Health (Luxembourg) by crossing *IL-22^ind^* with *Cd4^Cre^-Gclc^fl/fl^* mice.

*Cd4Cre IL-17A^ind^* mice were obtained by crossing *IL-17A^ind^* mice with *Cd4Cre* mice purchased from The Jackson Laboratory*. Cd4Cre Gclc^fl/fl^IL-17A^ind^* mice were generated at the Luxembourg Institute of Health (Luxembourg) by crossing *IL-17A^ind^* with *Cd4Cre Gclc^fl/fl^* mice.

All experiments used sex- and age-matched mice (8-12 weeks old) and were carried out using littermate controls. All animal experimentation protocols were approved and conducted according to the LIH Animal Welfare Structure guidelines.

### *Citrobacter rodentium* culture and infection

The nalidixic acid-resistant *Citrobacter rodentium* strain DBS100 was kindly provided by Dana Philipott (Toronto, Canada). Bacteria were suspended in 2.5% sterilized Miller’s Luria Bertani (LB) Broth (VWR) containing 75µg/mL nalidixic acid (Sigma-Aldrich) and cultured in a shaking incubator at 37°C with 160rpm rotation. When bacterial density achieved 0.6 OD^600^, mice were infected with 10^8^ CFU of *C. rodentium* in 100µL by oral gavage. The same bacterial culture was used to gavage all mice used in the same experiment. Food was withdrawn from mouse cages 16 hr prior to infection.

For *in vivo* administration of NAC (Sigma-Aldrich), experimental mice were supplied with drinking water containing 40 mM NAC whereas control groups received regular water. NAC administration commenced 7 days prior to infection, and fresh NAC-containing drinking water was provided every 3 days over the whole course of the infection.

For *in vivo* administration of IL-22, mice were injected intravenously (i.v.) with 100µg/100µL/mouse of either IL-22-Fc fusion protein (Genentech) or an isotype-matched IgG2A control-Fc protein (Ragweed:9652 10D9.W.STABLE mIgG2a; Genentech). Injections commenced one day before *C. rodentium* infection (day −1) and were repeated on days 3, 6, 9, 12 and 15 p.i.

### *Citrobacter rodentium* quantification in mouse feces, organs and blood

To determine CFUs of *C. rodentium* in feces and organs, fresh samples of feces and organs were harvested from infected mice at various time points (as indicated in the Figure). Samples were weighed and homogenized in 1mL cold PBS by continuous vortexing (feces) or with the help of a syringe plunger (organs). To measure CFUs of *C. rodentium* in blood, blood was withdrawn by cardiac puncture at the desired time points and collected in BD Microtainer® tubes (BECTON DICKINSON; 365986). Blood samples and homogenates of feces and organs were plated in serial dilutions (up to 10^9^) on LB agar composed of 2.5% sterilized LB Broth (Miller) (VWR) plus 1.5% of agar (Bioscience) containing 75µg/mL nalidixic acid (Sigma-Aldrich). Plates were incubated overnight at 37°C in a bacterial incubator. Numbers of CFUs in feces, organs and blood were determined by blinded counting of bacterial colonies. Results were normalized to the dilution and weight of the organ or fecal pellets or the volume of blood used.

### Fluorescein isothiocyanate (FITC)-Dextran gut permeability assay

To assess gut permeability, mice were orally gavaged with 150 μL of an 80 mg/mL solution of 3-5 kDa-FITC-Dextran (Sigma-Aldrich) in PBS. Mice were sacrificed at 4 hr post-gavage, and blood was collected in BD Microtainer® tubes (BECTON DICKINSON; 365986) and centrifuged at 5000 rpm for 10 min at 4°C. Plasma was diluted 5-fold and FITC-Dextran was quantified using a Mithras LB 940 (Berthold Technologies) fluorometer at an excitation wavelength of 485 nm and an emission wavelength of 535. Food was withdrawn from mouse cages 16 hr prior to FITC-Dextran administration.

### Colonic tissue histology and histochemistry

For histological analysis of mouse colons, distal colons containing fecal content were isolated at day 12 p.i. and fixed for 3 hr at room temperature (RT) in freshly prepared Carnoy’s fixative [60% anhydrous methanol (Sigma-Aldrich), 30% chloroform (Sigma-Aldrich), 10% glacial acetic acid (Sigma-Aldrich)]. Fixed colons were transferred to fresh Carnoy’s fixative and fixed again overnight. Fixed samples were washed in anhydrous methanol for 2 hr followed by transfer to fresh methanol and storage at 4°C until further use. Using a Tissue-Tek VIP processor (SAKURA), samples were subjected to consecutive washes with 100% denatured ethanol

(VWR), treatment with the clearing agent toluene (VWR), and paraffin embedment (Leica). Sections (3µm) were cut using a Rotary Microtome Microm HM 340E (Thermo Fisher Scientific). Paraffin-embedded sections were either stained with hematoxylin (Medite) plus eosin (VWR) (H&E), or Alcian Blue (Dako). Alcian Blue staining was performed using an Artisan Link Pro Special Staining System (Dako). Slides were scanned with an automated digital slide creation and viewing system (Philips IntelliSite Pathology Solution; 760001).

### Immunofluorescence staining and microscopy

To visualize tight junction localization and epithelial mucus content, distal colons were isolated on day 12 p.i. and feces were flushed out with cold PBS. Colons were cut open longitudinally and embedded flat, with the lumen facing downwards, in Tissue-Tek O.C.T. compound (SAKURA; 4583) in cryomolds and snap-frozen in liquid nitrogen. Cryosections (5µm) were prepared using a CM1850 UV Cryostat (Leica Biosystems) at −18°C and mounted on glass slides. For immunofluorescent analysis of colonic crypts, sections on glass slides were prepared as previously described (Lee et al., 2015). Briefly, samples were dried for 2 hr and fixed in 100% absolute ethanol (VWR) for 30 min at 4°C. Fixed samples were transferred to 100% acetone (cooled at -20°C) (Sigma-Aldrich) and incubated for 3 min at RT, followed by blocking with PBS containing 10% FBS (Biochrom GmbH) for 30 min at RT and washing twice in PBS. Primary antibodies recognizing either claudin-2 (Abcam) or MUC2 (Abcam) were diluted 1:200 in PBS+10% FBS and incubated with slides overnight at 4°C. After two washes in PBS, samples were stained overnight at 4°C with secondary goat anti-rabbit IgG-Alexa Fluor 594 antibody (Abcam) and Alexa Fluor 488 Phalloidin (Thermo Fisher Scientific). Washing was repeated and nuclei were stained with DAPI Fluoromount-G® medium (SouthernBiotech; 0100-20) for 24 hr at RT. Stained sections were visualized using a ZEISS Axio Observer microscope, and images were analyzed with ZEISS ZEN Blue 3.0 software. Colonic lumen diameters (for claudin-2 staining) were calculated using Fiji imageJ.

### Preparation of lamina propria of colon and caecum

Lamina propria of colon or caecum was isolated using the Lamina Propria Dissociation Kit (Miltenyi) following the manufacturer’s protocol. Briefly, colon or caecum tissues were resected and cut open longitudinally, and residual fat tissue was removed. Tissues were washed in HBSS without Ca^2+^, Mg^2+^ (Westburg) containing 10mM Hepes (Sigma-Aldrich).

Washed tissues were cut into small pieces and predigested in HBSS without Ca^2+^, Mg^2+^ containing 10mM Hepes, 5mM EDTA (Sigma-Aldrich), 5% FBS (Biochrom GmbH), 1mM dithiothreitol (DTT) (Sigma-Aldrich) for 20 min at 37°C using a rotator mixer (Intelli-Mixer, ELMI). Tissues were recovered in a 100µm strainer and re-incubated in fresh predigestion solution. After collection in a 100µm strainer, tissues were washed in HBSS without Ca^2+^, Mg^2+^ containing 10mM HEPES for 20 min at 37°C under continuous rotation. Again, tissues were recovered in a 100µm strainer and transferred into a C Tube (Miltenyi Biotec; 130-096-334) containing HBSS with Ca^2+^, Mg^2+^ (Westburg) and an enzyme mix (enzyme D, enzyme R, enzyme A) prepared from the Lamina Propria Dissociation Kit (Miltenyi) components. C Tubes were transferred into the GentleMACS Octo Dissociator with Heaters (Miltenyi Biotec; # 130-096-427) and the “37C_m_LPDK_1” program was run. Cold PBS containing 0.5% BSA (Sigma-Aldrich) was added to stop the reaction. Samples were passed through 40µm strainers, pelleted at 300xg for 10 min at 4°C, and resuspended in medium consisting of RPMI 1640 (Westburg) supplemented with 10% FBS (Biochrom GmbH), 1% penicillin/streptomycin (Gibco), 1% L-glutamine (Westburg), and 55µM 2-mercaptoethanol (Gibco). Lamina propria lymphocytes were analyzed by flow cytometry (see below).

### Naïve T cell isolation and *in vitro-*differentiation

Naïve CD4^+^ T cells were isolated from mouse spleen and lymph nodes by magnetic bead sorting using the Naïve CD4^+^ T cell isolation kit (Miltenyi Biotec) following the manufacturer’s protocol. Negative magnetic bead sorting was performed using the autoMACS® pro Separator (Miltenyi Biotec). Cell numbers were determined using a CASY cell counter (Omni Life Science). To induce *in vitro* differentiation, naïve T cells were cultured at 2×10^6^ cells/mL for 3 days in medium consisting of IMDM (Westburg) supplemented with 10% FBS (Biochrom GmbH), 1% penicillin/streptomycin (Gibco) and 55μM 2-mercaptoethanol (Gibco), and in the presence of Th cell subtype-specific cytokine mixes (see below).

For the induction of conventional Th17 cells, naïve T cells were cultured in the presence of TGF-β (2ng/µL; Bio-Techne), IL-6 (30ng/mL; Miltenyi Biotec), anti-IFN-γ (5µg/mL; BD Biosciences), soluble anti-CD28 (1µg/mL; Biolegend) and plate-bound anti-CD3 antibody (5 μg/mL; Biolegend). For the induction of IL-22-expressing Th17 cells, naïve T cells were cultured in the presence of IL-6 (30ng/mL; Miltenyi Biotec), soluble anti-CD28 (1µg/mL; Biolegend), plate-bound anti-CD3 antibody (5μg/mL; Biolegend or BD), IL-1β (50ng/mL; Miltenyi Biotec) and IL-23 (20ng/mL; Miltenyi Biotec).

For NAC experiments, the 3-day incubation period of *in vitro* T cell differentiation was conducted in culture medium containing 10mM of the antioxidant N-acetyl-cysteine (Sigma-Aldrich). For PI3K inhibitor treatment, 2 or 20 μM LY294002 or DMSO for control was added to culture medium for last 24h of the differentiation.

### DNA and RNA extractions

DNA from frozen fecal samples was isolated using the NucleoSpin DNA Stool kit (Macherey-Nagel) following manufacturer’s protocol.

For isolation of RNA from colon and caecum, tissues were collected in 1mL phenol-based TRIZOL (Thermo Fisher Scientific) and homogenized in a TissueLyser II (Qiagen) using Stainless Steel Beads (Qiagen; 69989). Samples were incubated at RT for 5 min, followed by addition of 0.2 mL chloroform (Sigma-Aldrich) per sample. Samples were shaken for 15 seconds, incubated for 3 min at RT, and centrifuged at 12,000xg for 15 min at 4°C. The aqueous phase containing the RNA was collected and 0.5 mL isopropanol (VWR) was added per sample. Samples were transferred onto NucleoSpin RNA columns (Macherey-Nagel) to purify RNA using the NucleoSpin RNA Kit protocol. Briefly, after binding to columns, RNA was treated with DNase, washed and eluted in RNase-free water.

For isolation of RNA from *in vitro* cultured Th17 cells, NucleoSpin RNA XS kit (Macherey-Nagel) was used according to protocol.

DNA and RNA concentrations were quantified using a NanoDrop 2000c Spectrophotometer (Thermo Fisher Scientific).

### Real-time reverse transcription polymerase chain reaction (RT-qPCR)

For qPCR of stool DNA samples, 6μL DNA (50 ng) was mixed with 10µL SYBR™ Fast SYBR™ Green Master Mix (FISHER SCIENTIFIC), 2pmol of forward primer and 2pmol of reverse primer (please see list of primers in Reagents). Reactions were run on a ABI 7500HT Fast qRT-PCR instrument.

For RT-qPCR of RNA samples, 2μL RNA (150 ng) was mixed with 5μL Master Mix (Luna Universal One-Step RT-qPCR Kits; Bioké), 0.3μL reverse transcriptase, 2.7μL H^2^O, 2pmol of forward primer and 2pmol of reverse primer (cf. Reagents). Reactions were run on a CFX384 instrument (Bio-Rad). Transcript data were normalized to total Eubacteria (for bacterial DNA in feces), rps17 (for bacterial virulence factors), and hprt (for antimicrobial peptides), and analyzed using the ΔΔCt method as previously described (Mak et al., 2017).

### Flow cytometry

Flow cytometric staining and analyses were performed as previously described (Cossarizza et al., 2019). T cells were identified as either CD3^+^CD4^+^ or CD4^+^ alone (see Figure Legends). LTi cells were identified as Lineage^-^ [i.e. (Ter119, CD19, CD11b, CD5, Ly6G/Ly6C)^-^] CD3^-^Nkp46^-^ CD4^+^.

To stain surface molecules, cells were incubated in FACS buffer (PBS containing 1% FBS and 5mM EDTA, pH 8.0) in the presence of antibodies (cf. Reagents; section Antibodies). Antibodies were diluted 1:200 and incubated for at least 30 min at 4°C, protected from light. Stained cells were washed in FACS buffer prior to flow cytometric analysis.

For intracellular staining to detect p-mTOR, p-PI3K and p-AKT(T308), cells were either fixed and permeabilized using BD Cytofix/Cytoperm Fixation/Permeabilization kit (BD) or fixed in 2% formaldehyde (Sigma-Aldrich) for 10 min at RT, permeabilized in 0.01% saponin (Sigma-Aldrich), and incubated for 30 min with antibody (diluted 1:200 in saponin) at 4°C in the dark. Stained cells were washed and resuspended in saponin. For intracellular staining of p-4E-BP1 (Thr37/46), cells were fixed with 4% formaldehyde (Sigma-Aldrich) for 15 min at RT, resuspended in 10 ul of PBS and permeabilized with ice-cold MetOH (Sigma-Aldrich) for 20 min on ice, protected from light, washed 3x in PBS and stained in FACS buffer with antibodies for 1h at 4°C and washed in FACS buffer before flow cytometry analysis. For intracellular staining of transcription factors (including RORγT, Tbet), cells were fixed for 40 min at 4°C using the eBioscience™ Foxp3/Transcription Factor Fixation kit (Thermo Fisher Scientific) and permeabilized using the kit’s permeabilization buffer. Cells were incubated for 30 min with antibodies (diluted 1:200 in permeabilization buffer) at 4°C in the dark. Stained cells were washed in permeabilization buffer before flow cytometry analysis.

For intracellular staining of cytokines, cells were stimulated *in vitro* for 5 hr with phorbol 12-myristate 13-acetate (PMA; Sigma-Aldrich, 50ng/mL), calcium ionophore A23187 (Sigma-Aldrich, 750ng/mL), and BD GolgiPlug™ Protein Transport Inhibitor (BECTON DICKINSON, 1:1000 dilution). Stimulated cells were washed once in FACS buffer before fixation for 20 min at 4°C using the BD Cytofix/Cytoperm solution and permeabilization using BD Perm/Wash™ buffer (BD Biosciences). Permeabilized cells were incubated for 30 min with antibodies (diluted 1:200 in BD Perm/Wash) at 4°C in the dark. Stained cells were washed in permeabilization buffer before flow cytometry analysis.

To stain intracellular thiols, cells were processed as described previously (Franchina et al., 2022). Briefly, cells were stained at 37°C for 30 min in either complete RPMI 1640 (10% FBS, 1% penicillin/streptomycin, 1% L-glutamine, 55µM 2-mercaptoethanol) for lamina propria cells, or in complete IMDM (10% FBS, 1% penicillin/streptomycin, 55µM 2-mercaptoethanol) for *in vitro-*differentiated Th17 cells. Monobromobimane (mBBr) (Thermo Fisher Scientific) was added 10min before washing off the supernatant. Stained cells were washed twice and resuspended in cold PBS for flow cytometric aquisition.

For determination of intracellular or mitochondrial ROS levels, cells were stained at 37°C for 30 min with 2 µM 2’,7’-dichlorodihydrofluorescein diacetate (H^2^DCF-DA; Thermo Fisher Scientific) or 1 µM MitoSOX (Thermo Fisher Scientific), respectively. Cells were washed and resuspended as above. To quantify mitochondrial membrane potential or mass, cells were stained at 37°C for 30 min with 100nM MitoTracker Deep Red or 10nM MitoTracker Green (Thermo Fisher Scientific), respectively. Cells were washed and resuspended as above.

Dead cells were excluded from flow cytometric analyses by staining with either DAPI (Thermo Fisher Scientific), LIVE/DEAD® Fixable Near-IR dye (Biolgend), LIVE/DEAD® Fixable Green dye (Biolegend) or 7-AAD (Thermo Fisher Scientific). Flow cytometry was performed using a BD Fortessa instrument (BD Biosciences), and results were analyzed using FlowJo v10.6.2 software (Tree Star).

### ELISA measurement of cytokines and lipocalin-2

IL-17 or IL-22 levels in culture supernatants were quantified using the Mouse IL-17 DuoSet ELISA kit (Bio-Techne) or the Mouse IL-22 DuoSet ELISA kit (Bio-Techne), respectively. The Mouse Lipocalin-2/NGAL DuoSet ELISA kit (Bio-Techne) was used to determine lipocalin-2 concentrations in fecal supernatants obtained by homogenization of fecal pellets in PBS. All assays were performed following the manufacturer’s instructions.

Briefly, plates were coated with target-specific antibody overnight, washed in wash buffer, and blocked by incubating with reagent diluent for at least 1 hr. After washing, samples and the appropriate standards were added to plates and incubated for 2 hr. After washing, the corresponding biotinylated detection antibody was added for 2 hr. After washing, streptavidin coupled to horseradish peroxidase (HPR) was added for 20-30 min. Plates were washed and a substrate solution containing hydrogen peroxide was added. The HPR reaction was stopped by adding sulfuric acid. Optical density was determined using a Versa Max microplate reader (Molecular Devices) with SoftMax Pro7.1 software set to 450nm. Wavelengths were corrected by subtraction of background measurements at 570nm. Protein concentrations were determined and normalized to total cell numbers per culture (for IL-17 and IL-22) or fecal weight (for lipocalin-2).

### Luminescence assays

For quantification of GSH content in LP CD4^+^ cells, viable CD4^+^ cells were FACS-sorted using a FACSAria III (BD Biosciences) instrument according to published cell sorting guidelines (Cossarizza et al., 2019). Cells (1.5×10^4^/well) were subjected to a GSH/GSSG luminescence-based assay (Promega) following the manufacturer’s protocol. For quantification of ATP levels in *in vitro*-differentiated Th17 cells, cells (1×10^5^) were subjected to the CellTiter-Glo® assay (Promega) following the manufacturer’s protocol. Luminescence intensities were quantified using a Mithras LB 940 instrument (Berthold Technologies).

### Measurement of oxygen consumption rate (OCR) by Seahorse analysis

Seahorse analyses were performed using an XFe96 Extracellular Flux Analyzer (Agilent). Briefly, *in vitro-*differentiated Th17 cells were seeded in XF Seahorse DMEM medium (Agilent Technologies) supplemented with 1mM sodium pyruvate (Gibco), 2mM glutamine (Westburg), and 25mM glucose (Sigma-Aldrich) at a density of 2×10^5^ cells/well on a Seahorse XFe96 cell culture plate (Agilent Technologies; 101085-004). Plates were pre-coated with Corning™ Cell-Tak Cell and Tissue Adhesive (Thermo Fisher Scientific) containing 0.1M of sodium bicarbonate (Sigma-Aldrich). OCR was measured using the XF Cell Mito Stress Test (Agilent) according to manufacturer’s protocol. Sequential injections of 1µM oligomycin A (Sigma-Aldrich), 3µM carbonyl cyanide 4-(trifluoromethoxy)phenylhydrazone (FCCP) (Sigma-Aldrich), and 1µM antimycin A/rotenone (Sigma-Aldrich) were performed, with three measurements taken after each treatment.

Basal OCR was calculated from raw OCR values measured just before oligomycin injection. Maximal OCR was calculated from raw OCR values obtained from the second measurement after FCCP injection.

### Immunoblot analysis

For detection of claudin-2, claudin-3 and claudin-15 proteins, distal colons were cut open and washed in cold PBS. The tissue was cut into very small pieces using a clean razor blade and transferred into 0.3 mL RIPA buffer containing 50 mM Tris-HCl pH7-8 (Sigma-Aldrich), 150 mM NaCl (Sigma-Aldrich), 0.5% sodium deoxycholate (Sigma-Aldrich), 1% NP-40 (Abcam), 0.1% SDS (Carl Roth), 2mM EDTA (Sigma-Aldrich) and Protease/Phosphatase Inhibitor Cocktail (100x) (Bioké). Samples were incubated for 30 min on ice, then sonicated for 20 min, followed by another 30 min incubation on ice. Samples were centrifuged at 16,000xg at 4°C for 20 min and the clear supernatant was collected. This procedure was repeated. Lysates were diluted 1:3 in sample buffer containing 187.5 mM Tris-HCl pH 6.8, 6% SDS, 30% glycerol (Carl Roth), 0.03% bromophenol blue (Sigma-Aldrich) and 10% 2-mercaptoethanol. Samples were incubated for 5 min at 95°C, loaded on a Novex™ WedgeWell™ 16% Tris-Glycine gradient gel (Thermo Fisher Scientific; XP00162), and run at 100V for 90-100 min. Proteins were transferred onto a nitrocellulose membrane (iBlot2 Transfer Stacks, Nitrocellulose Mini; Fisher Scientific, 15239296) using an iBlot2 machine (Thermo Fisher Scientific) and blocked with 5% milk (Carl Roth) for 1hr. After washing with PBS-Tween (PBS-T), primary antibodies recognizing claudin-2 (1:1000; Abcam), claudin-3 (1:200; Thermo Fisher Scientific), claudin-15 (1:200; Thermo Fisher Scientific), or actin (1:5000; Sigma-Aldrich), which were diluted in PBS-T containing 5% BSA, were added to blots and incubated overnight at 4°C. Membranes were washed and secondary mouse anti-rabbit IgG-HRP (Santa Cruz Biotechnology), which was diluted 1:5000 in PBS-T with 5% milk, was added for 1 hr at RT. Proteins were visualized using Luminata™ Crescendo Western HRP substrate (Thermo Fisher Scientific) and an INTAS ECL Chemocam Imager.

### Bulk RNA sequencing & data analysis

RNA extraction of *in vitro* cultured Th17 cells was performed with the NucleoSpin RNA XS Kit (Macherey-Nagel) according to the manufacturer’s protocol. RNA concentrations and integrity were measured using a RNA 6000 NanoKit (Agilent) on a 2100 Bioanalyzer (Agilent).

Sequencing was performed by the Sequencing Platform of the Luxembourg Center for Systems Biomedicine (LCSB) of the University of Luxembourg. Samples were prepared using an Illumina Stranded mRNA library prep kit (Illumina) with the addition of IDT for Illumina DNA/RNA UD indexes (Illumina). Paired-end sequencing was executed using an Illumina NextSeq2000 machine with a read length of 2 x 50 bp.

RNA-seq transcript alignment was performed with Salmon (Patro et al., 2017) against the Mouse Transcriptome from Genecode release M30 assembly GRCm39 (Frankish et al., 2018). Subsequent analysis was conducted in R, and Tximeta (Love et al., 2020) was used to assign transcripts to genes before differential analysis with DESeq2 (Love et al., 2014). Gene set enrichment analysis (GSEA) was performed using ClusterProfiler (Yu et al., 2012).

### Bioinformatics analysis of RNA sequencing dataset GSE109142

The publicly available dataset GSE109142, which contains RNA sequencing TPM counts of rectal biopsies from pediatric ulcerative colitis (UC) patients (Haberman et al., 2019), was downloaded from the Gene Expression Omnibus (GEO) repository and processed using R version 4.1.0. Before applying the log2 transformation, for each gene of interest, half of the smallest non-zero value was added to the TPM counts. The lower and upper quartile values of *GCLC* expression were used to classify the patients according to low, intermediate and high expression of *GCLC*. The heat-map representation was generated using the ComplexHeatmap package (Gu et al., 2016) in R. The provided IBD MAYO scores (ranging from 0 to 12) were used to define disease severity as normal/mild (score < 6), moderate (score < 11) and severe (score

≥ 11) (Teixeira et al., 2015). The Th17 score was defined as the mean of scaled log2 gene expression values composing the signature (*IL-17A, IL-17F, IFN-γ, CD3E, CD4, STAT3, STAT5a, STAT1, STAT4, STAT6, AHR, RORC, RORA and TBX21*). To analyze the correlation between gene expression values or scores, the Pearson’s correlation coefficient (r) were calculated and significances (p-value) as well as r^2^ values shown.

### Quantification and statistical analysis

Data are expressed as the mean ± SEM with at least n=3 per group (refer to Figure Legends for detailed information). P values were determined by unpaired Student’s t test, one-way ANOVA or two-way ANOVA using Prism 8.0 (GraphPad). P values of ≤0.05 were considered statistically significant and are indicated with one or more asterisks (*p ≤ 0.05; ** p ≤ 0.01; ***p ≤ 0.001; ****p ≤ 0.0001; ns: not significant).

## Reagents

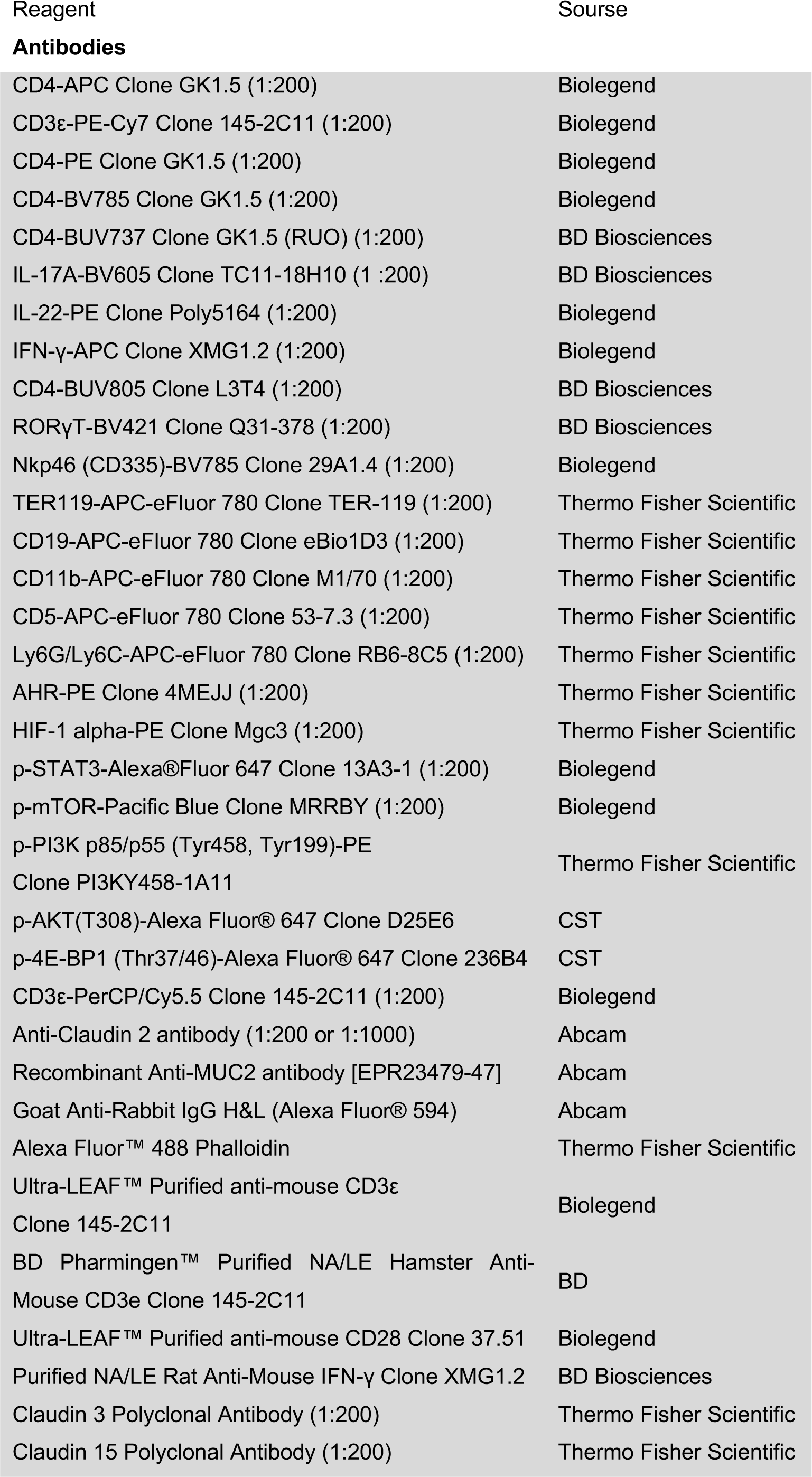
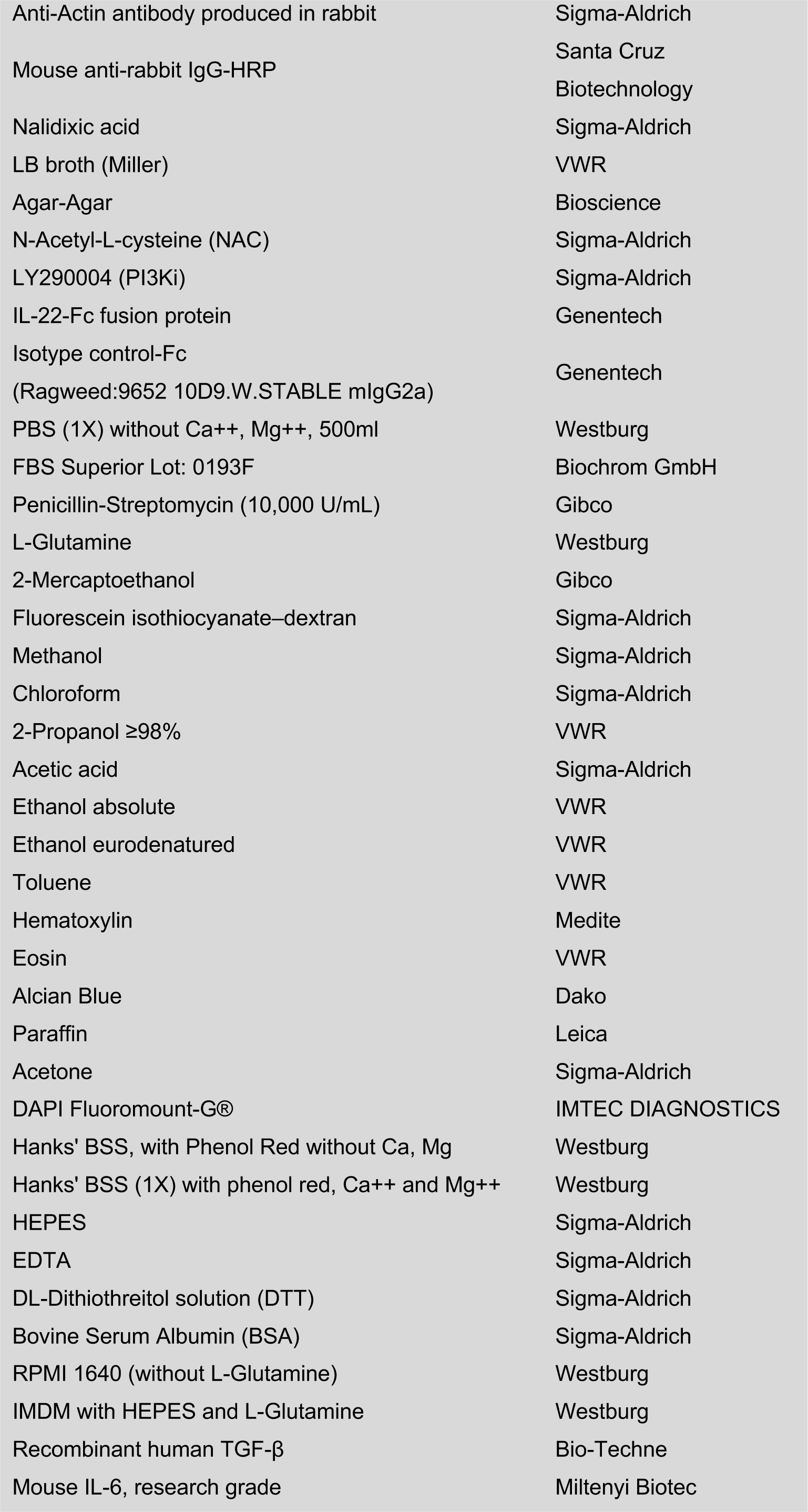
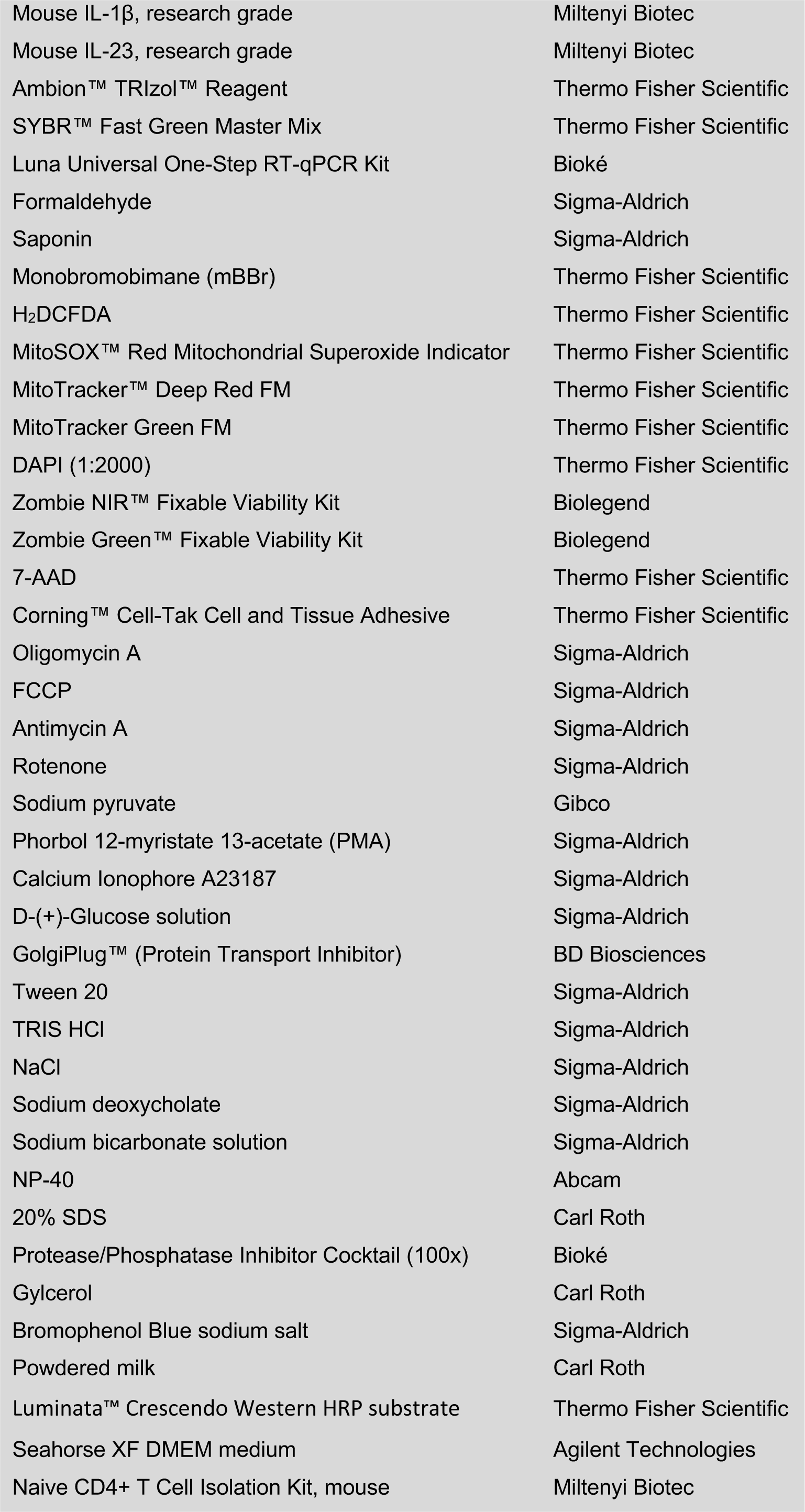
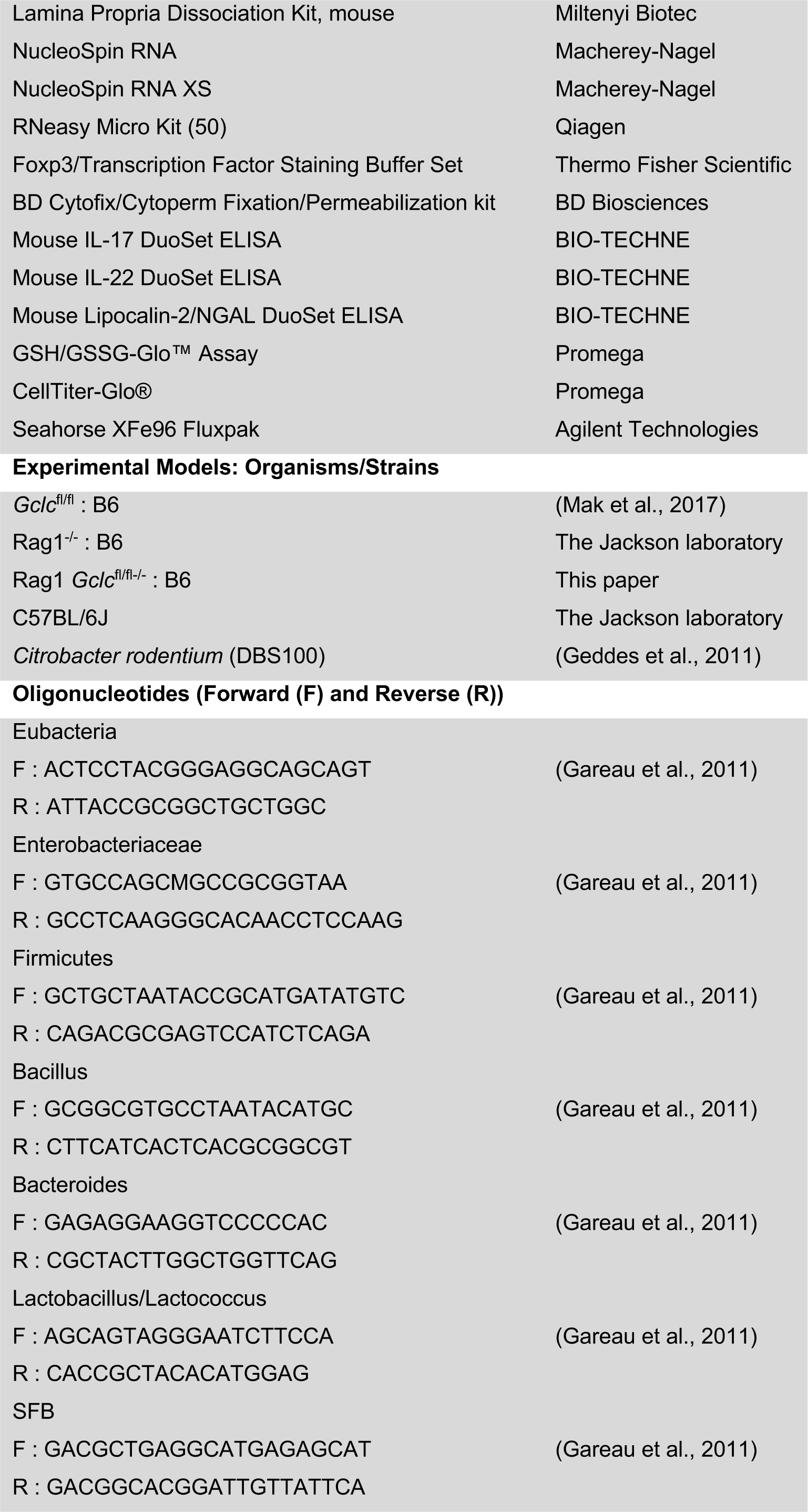
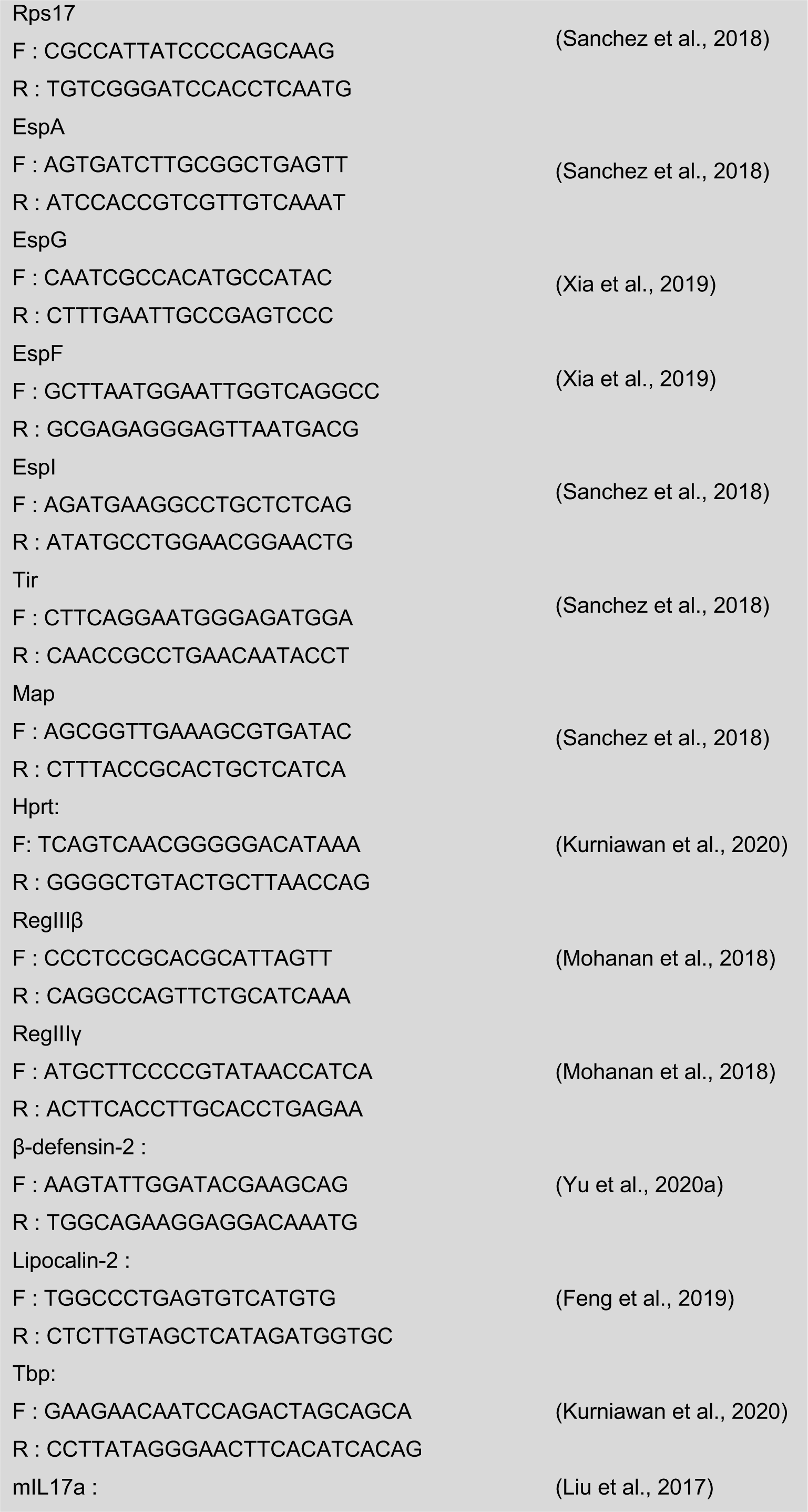
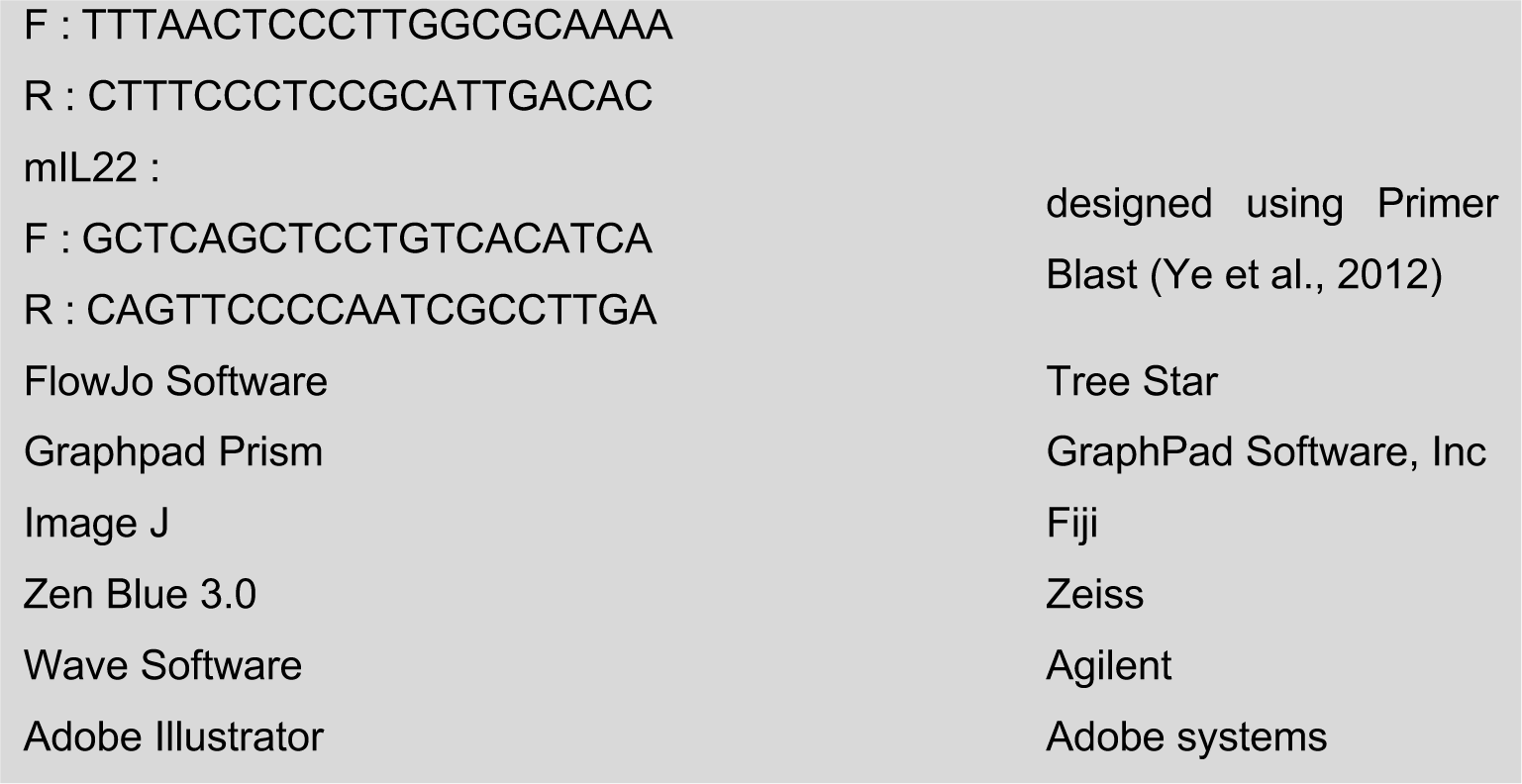

## Notes

### Competing Interest Statement

The authors have declared no competing interest.

